# Disruption of Microhomology-mediated End-joining in Ewing Sarcoma

**DOI:** 10.1101/2025.05.06.651696

**Authors:** Shuhei Asada, Guangli Zhu, Jithma Prasad Abeykoon, Yutaro Tanaka, Huy Nguyen, Yuna Hirohashi, Divya R. Iyer, Nicholas William Ashton, Sirisha Mukkavalli, Martha Velazquez, Lige Jiang, Miles Del Busso, Judith Jebastin Thangaiah, Steven I. Robinson, Kalindi Parmar, Eliezer M. Van Allen, Riaz Gillani, Geoffrey I. Shapiro, Alan D. D’Andrea

## Abstract

Ewing sarcoma (EwS) is a group of bone and soft tissue cancers in children and young adults. Since EwS cells have pronounced sensitivity to radiation and chemotherapy-induced DNA damage, the role of the oncoprotein, EWS-FLI1, in DNA repair is likely. Here, we demonstrate that EWS-FLI1 causes a defect in microhomology-mediated end-joining (MMEJ) repair. EWSR1 is a splicing factor that promotes the faithful splicing of the *POLQ* pre-mRNA, required for the expression of POLΘ, a critical protein in the MMEJ pathway. Expression of EWS-FLI1, or loss of EWSR1, causes exon 25 skipping of the *POLQ* transcript, decreased POLΘ expression, impaired MMEJ, and cellular sensitivity to inhibitors of the Fanconi Anemia (FA), NHEJ, or HR pathways, through the mechanism of synthetic lethality. Knockdown of EWS-FLI1 expression restores POL0 mitotic foci and increases MMEJ activity. Inhibitors of the FA, NHEJ, or HR therefore may provide a targeted therapy for patients with EwS.

**Highlights:**

- Ewing sarcoma tumors have a deficiency in POLθ expression and a corresponding loss of MMEJ activity
- EWSR1 is a splicing factor that interacts with other splicing factors such as FUBP1 and KHSRP/FUBP2 to accurately splice the *POLQ* mRNA.
- The EWS-FLI1 fusion oncoprotein, or loss of EWSR1, causes a splicing defect, leading to exon25 skipping of the *POLQ* pre-mRNA and loss of POLθ expression
- The MMEJ deficiency of EwS cells results in cellular sensitivity to inhibitors of Fanconi Anemia, Homologous Recombination or Non-Homologous End-Joining
- Exon 25 skipping of *POLQ* mRNA is a predictive biomarker for HR inhibitors in human cancers

## INTRODUCTION

Ewing sarcoma (EwS) is a group of bone and soft tissue cancers in children and young adults^1,2^. EwS tumor cells have a chromosome 11;22 translocation, resulting in expression of the EWS-FLI1 fusion oncoprotein. Their elevated sensitivity to ionizing radiation and DNA-damaging chemotherapy, as well as their relatively low level of somatic mutations, suggest that EwS cells have an underlying defect in an error-prone DNA repair (DDR) pathway. Despite this high drug sensitivity, novel treatments for EwS are required, especially for patients with relapsed and/or refractory disease, who have a low 5-year overall survival rate of only 10%.

Recent studies have identified some abnormalities in DDR in Ewing sarcoma cells. Expression of EWS-FLI1 fusion protein promotes R-loop formation and impaired BRCA1 recruitment to chromatin, suggesting a defect in the homologous-recombination repair pathway (HRD) ^3^. Despite this preclinical observation, PARP inhibition, an approved treatment for HR-deficient tumors, was ineffective in the treatment of EwS patients^4^. Other cellular defects resulting from EWS-FLI1 expression may indirectly affect DDR, such as defects in BAF complex-mediated transcription ^5,6^, condensate generation ^7–9^, and mRNA splicing ^10^. However, the mechanism by which EWS-FLI1 specifically influences DDR pathways is unknown, and other more profound DNA repair defects, beyond the HRD phenotype, may exist.

DNA double-strand breaks (DSBs) are the most critical forms of DNA damage in cells. There are three main pathways for repairing DSBs -non-homologous end-joining (NHEJ), homologous recombination (HR), and microhomology-mediated end-joining (MMEJ). For a cancer cell to survive and proliferate, DSBs must be repaired, and the cell must rely on at least one of these pathways. DNA polymerase theta (POL0), an error-prone polymerase encoded by the *POLQ* gene, is required for the MMEJ pathway. Overexpression of *POLQ* confers chemoresistance, whereas depletion of POLΘ results in cancer predisposition and hypersensitivity to irradiation and chemotherapy ^11,12^. *To date, there are no reported human cancers with defects in the MMEJ pathway*.

Here, we demonstrate that EwS cells have an underlying defect in MMEJ. Expression of EWS-FLI1, or loss of EWSR1, in these cells promotes exon skipping of the *POLQ* pre-mRNA, generating a premature termination codon and a profound reduction of POLΘ protein expression. EWS-FLI1 expressing cells exhibit a phenotype like *POLQ* deficient cells, including hypersensitivity to topoisomerase inhibitors, irradiation, and synthetic lethality with loss of the Fanconi Anemia, NHEJ, or HR pathways. Knockout of the EWS-FLI1 oncoprotein in EwS cells results in reduced exon skipping, re-expression of the POLΘ protein, and a rescue of MMEJ activity. Accordingly, the DDR defect in EwS tumors could be exploited for therapeutic benefit by inhibiting another DSB repair pathway.

## RESULTS

### *EWSR1* knock-out cells phenocopy MMEJ-deficient cells

Previous CRISPR screens and Depmap analyses have demonstrated that genes in the MMEJ pathway are synthetic lethal in cells with an underlying defect in an HR gene ^13,14^, Interestingly, in these screens, knockout of the *EWSR1* gene was highly similar to knockout of known MMEJ genes, including *POLQ* ^15^, *XRCC1* ^16^, *APEX2* ^17^, and *RHNO1* ^18^, in the killing of HR-deficient tumor cells (**Fig. S1A)**. To investigate the function of EWSR1 in DNA repair, we initially generated an *EWSR1* knock-out in RPE *TP53-/-* cells (**Fig. 1A**). *EWSR1* knockout resulted in hypersensitivity to ionizing irradiation, topoisomerase inhibitors, and ATR inhibitors, but not to the PARP inhibitor, suggesting a loss of MMEJ activity (**Fig. S1 B-F**). The DDR defect observed in *EWSR1* knockout cells was similar to the defect observed in *POLQ* knockout cells ^11^. Consistently, *EWSR1* is mapped closely to *POLQ* in the previous repair-seq, indicating a functional relationship between these genes in DNA damage repair ^19^.

**Figure 1:**
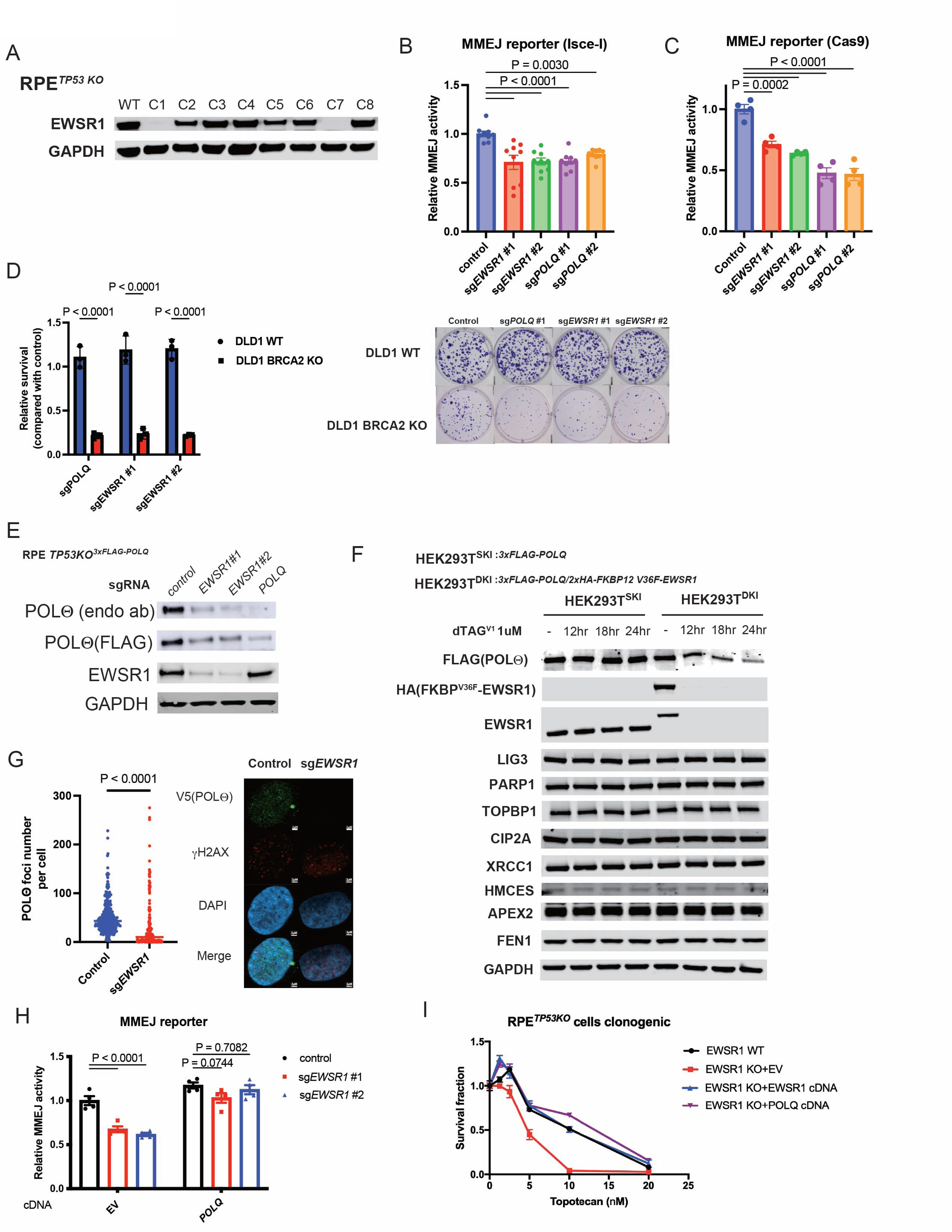
*EWSR1* knock-out cells display MMEJ-deficient phenotypes through POL0 protein reduction. (A) Immunoblotting analysis for the detection of EWSR1 in RPE p53-/-clones edited with CRISPR-Cas9. (B, C) U2OS I-SceI-based (B, n=9) or Cas9-based (C, n=4) or were transduced lentivirally transduced with control sgRNA or sgRNA targeting *EWSR1* or *POLQ* (co-expressing puromycin resistance gene). Cells were treated with puromycin for 4 days, and then subjected to MMEJ reporter analysis. (D) Colony formation assay data of DLD1 *BRCA2* wild-type and *BRCA2* knockout cells transduced with control, sg*POLQ* or sg*EWSR1* (n=3, left). Shown are representative images of the colonies (right). (E) RPE *TP53-/-* 3xFLAG-*POLQ* knock-in cells were lentivirally transduced with Cas9 and a sgRNA targeting control (NT), *EWSR1* or *POLQ* (co-expressing puromycin resistance gene). After puromycin selection, cells were subjected to immunoblotting analysis. (F) HEK293T 3xFLAG-*POLQ* single knock-in cells and HEK293T 3xFLAG-*POLQ*/2xHA-FKBP12 V36F *EWSR1* double knock-in cells are treated with DMSO or dTAG^V1^ for indicated time. Cell lysates were extracted and followed by immunoblotting analysis. (G) Immunofluorescence analysis showing DNA damage-induced the foci formation of V5-tagged endogenous POL0 was decreased by EWSR1 depletion (scale bars = 2μm). (H) U2OS MMEJ reporter cells (I-SceI based) were transduced with a control vector or POLQ cDNA (co-expressing blasticidin resistance gene). After blasticidin selection, cells were subsequently transduced with Cas9 and a sgRNA targeting control or EWSR1 (co-expressing puromycin resistance gene). After puromycin selection, cells were subjected to MMEJ reporter analysis (n=4). (I) RPE *p53*-/-cells and RPE *p53*-/-*EWSR1* knockout cells were transduced with control, wildtype *EWSR1* or *POLQ* cDNA. Colony formation assays were performed in the presence of Topotecan (n=3).

Next, the impact of *EWSR1* loss on the activity of DNA double-strand break repair pathways in U2OS cells was evaluated, using established Isce-I-based reporter systems ^20,21^. *EWSR1* knockout mildly decreased HR activity (**Fig. S1G**); however, the *EWSR1*-deficient cells were not sensitive to the PARP inhibitor, Olaparib. Interestingly, *EWSR1* knockout significantly decreased MMEJ activity (**Fig. 1B**). To further confirm a MMEJ reduction upon *EWSR1* depletion, a novel Cas9-based MMEJ reporter system was generated (**Fig. S1H**). Again, *EWSR1* knockout decreased MMEJ activity in this Cas9-based reporter cells (**Fig. 1C**). Consistent with this result, *EWSR1*-deficient cells were resistant to the *POLQ* inhibitor, Novobiocin ^22^(**Fig. S1I**).

Recent studies suggest that *EWSR1* knockdown is synthetic lethal with BRCA2 loss ^13,14^. To validate these results, depletion of EWSR1 in BRCA2-proficient and BRCA2-deficient DLD1 cells was evaluated. Like *POLQ* loss, EWSR1 loss profoundly decreased colony-forming activity in the *BRCA2*-deficient DLD1 cells (**Fig. 1D**). EWSR1 loss also efficiently decreased clonogenic activity of the *BRCA1*-mutated breast cancer cell lines, MDA-MB-468 and HCC1937 cells (**Fig S1. J, K**). To further confirm the synthetic lethality of EWSR1 and BRCA1/2, we generated RPE *TP53-/-* cells stably expressing Cas9 (**Fig. S1L)**. These cells were subsequently lentivirally-transduced with a sgRNA targeting control, *POLQ,* or *EWSR1* (co-expressing GFP) together with another sgRNA targeting control, *BRCA1* or *BRCA2* (co-expressing tRFP657), and changes in the frequency of GFP/tRFP657 cells were monitored. Again, like *POLQ* loss, *EWSR1* loss resulted in synthetic lethality with *BRCA1/2* loss in this system. Taken together, *EWSR1* functions as a MMEJ gene.

### EWSR1 promotes MMEJ by maintaining POL0 expression

Since EWSR1 loss caused a reduction of MMEJ activity, we examined the impact of EWSR1 loss on protein expression of known MMEJ-related proteins. To monitor the POL0 protein expression, RPE *TP53-/-* cells expressing 3xFLAG-tagged *POLQ* were generated. Interestingly, *EWSR1* knockout resulted in decreased POLΘ expression (**Fig. 1E, Fig. S2A**). Increased radiosensitivity of the *EWSR1* knockout cells was observed (**Fig. S2B**). To further evaluate the direct and acute effect of EWSR1 loss on POLΘ expression, we tagged endogenous EWSR1 protein with the mutant cytosolic prolyl isomerase FKBP12^F36V^ in endogenously 3xFLAG-tagged *POLQ* expressing HEK293T cells ^23,24^. In this targeted protein degradation system, the small bifunctional molecule dTAG^v1^ bridges FKBP12^F36V^ with the E3 ubiquitin ligase VHL, resulting in substrate ubiquitination and proteasomal degradation. Treatment of cells harboring the FKBP12^F36V^-tagged EWSR1 with dTAG^v1^ resulted in rapid degradation of EWSR1 protein (**Fig. S2C-F**). Interestingly, this acute depletion of EWSR1 rapidly reduced the expression of POL0 but did not affect the expression of other known MMEJ-related proteins, including APEX2, HMCES, FEN1, LIG3, PARP1, and XRCC1 nor HR-related proteins such as BRCA1 and BRCA2 (**Fig 1F, Fig. S2G).** Since EWSR1, as well as FUS and TAF15, belongs to the FET family of proteins ^25^, we speculated that POL0 loss and MMEJ deficiency might be a common phenotype of the loss of FET family genes. However, FUS or TAF15 loss did not affect the protein level of POL0 or MMEJ activity (**Fig. S2H, I**). Consistently, *EWSR1* knockout reduced the POL0 foci assembly upon irradiation-induced DNA damage (**Fig. 1G**).

The functional significance of the loss of POL0 on MMEJ activity in *EWSR1*-depleted cells was next examined. Transduction with the WT *POLQ* cDNA rescued MMEJ activity in *EWSR1* knockout cells (**Fig 1H**). Moreover, *POLQ* cDNA transduction corrected the hypersensitivity to the topoisomerase inhibitors, indicating that loss of POL0 protein is responsible for these phenotypes in *EWSR1* knockout cells (**Fig 1I, Fig S2J**). Like POL0 loss, EWSR1 loss increased unrepaired DNA damage and micronuclei in BRCA-deficient cells (**Fig S2K, L**). Collectively, EWSR1 promotes MMEJ via the maintenance of POL0 protein expression.

### EWSR1 directly binds to the *POLQ* transcript and prevents *POLQ* exon skipping

To investigate the cause of the reduced POL0 protein expression following loss of EWSR1, we evaluated the stability of POL0 protein in the presence or absence of EWSR1. U2OS MMEJ-reporter cells were transduced with the *POLQ* cDNA, followed by *EWSR1* knockout by the CRISPR-Cas9 system. However, the protein level of POL0 expressed by *POLQ* cDNA did not change after *EWSR1* depletion (**Fig. S3A**). Furthermore, a proteasome inhibitor MG132, or the autophagy inhibitor chloroquine, did not rescue endogenous POL0 expression in *EWSR1*-depleted cell (**Fig. S3B**), suggesting a pre-translational mechanism.

To assess the overall influence of EWSR1 loss on transcript levels, including the POLQ transcripts, we next performed an RNA-seq analysis on RPE1 *TP53-/-* cells and RPE1 *TP53-/-EWSR1* knockout cells. Alternative splicing events were analyzed by the MISO framework ^26^ as well as the rMATs framework^27^. Based on this analysis, various alternative splicing events, including exon skipping events, were observed in *EWSR1* knock-out cells (**Fig. S3C-G)**. Among DNA repair genes with exon skipping events, we identified several exon skipping events of the *POLQ* transcript, including exon 7, exon 21 and exon 25, as top hits (**Fig. 2A**). Among these exon skipping events within the *POLQ* transcript, the skipping of exon 25 was the most prominent (**Fig. 2B)**. Of note, *POLQ* exon 25 is located between long introns, intron 24 and intron 25. Furthermore, we re-analyzed previously published RNA-seq datasets using si-control and si-EWSR1 cells, and again the *POLQ* exon 25 skipping is one of the most significant events (**Fig. S3H, I**). We validated the results in multiple *EWSR1* knockout clones by RT-PCR (**Fig. S3J**).

**Figure 2:**
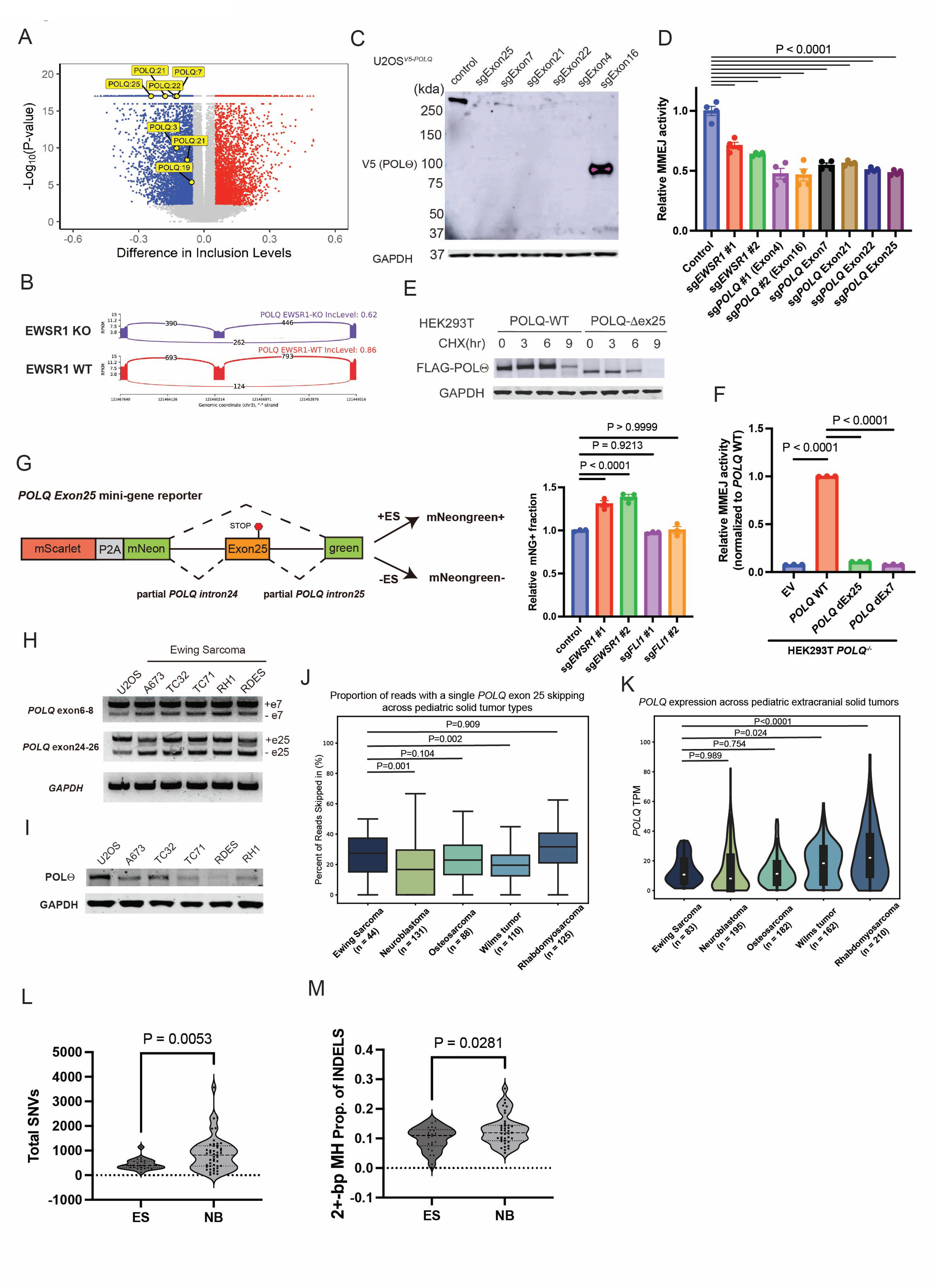
Disruption of MMEJ activity in Ewing sarcoma is driven by *POLQ* exon skipping. (A) Volcano plots showing alternatively spliced skipped exon (SE) in RPE *p53*-/-*EWSR1* knock-out cells compared with RPE *p53*-/-cells defined by deep RNA-seq analysis. (B) Sashimi plots of the alternative splicing events in control and *EWSR1* knock-out RPE *p53*-/-cells: the region between exon24 and exon26 of the *POLQ* transcript. (C) U2OS endogenous V5 tagged-POLΘ expressing cells were transduced with Cas9 and sgRNA targeting the indicated exon of the *POLQ* gene. Cell lysates were extracted and followed by immunoblotting analysis. (D) U2OS MMEJ ver.2 (Cas9-based) cells were lentivirally transduced with control sgRNA or sgRNA targeting different *POLQ* exons (co-expressing puromycin resistance gene). Cells were treated with puromycin for 4 days, and then subjected to MMEJ reporter analysis (n=4). (E) *POLQ* mRNA without exon 25 produces shorter POLΘ which is unstable. Blot showing HEK293T cells transfected with FLAG-*POLQ* cDNA containing *POLQ* full-length CDS or CDS lacking exon25. Cells were treated with CHX for 3–9 hr. (F) HEK293T *POLQ* knockout cells were transfected with MMEJ reporter (ver.2) and sgRNA targeting the reporter together with control vector or indicated *POLQ* cDNA. 48h after transfection, cells were subjected to FACS analysis to measure the MMEJ activity (n=3). (G) Schematic representation of the minigene reporter construct (left). K562 cells carrying the minigene reporter were lentivirally transduced with Cas9 and sgRNA targeting the indicated genes. Shown are relative frequencies of mNeonGreen positive fraction within mScarlet positive cells (right, n=3). (H) RT-PCR showing the exon 7 and exon 25 skippings of *POLQ* transcript in Ewing sarcoma cell lines. (I) Immunoblotting analysis showing the lower levels of POL0 expression in Ewing sarcoma cell lines compared to U2OS cells. (J) rMATS analysis showing *POLQ* exon skipping events in primary samples from pediatric cancer patients. n = 498 (EWS = 44, NBL = 131, OS = 88, WT = 110, RMS = 125). (K) RNA-seq data from the patient sample shows the *POLQ* mRNA expression is significantly lower in Ewing sarcoma patients compared with RMS/WT. n = 832 (EWS = 83, NBL = 195, OS = 182, WT = 162, RMS = 210). (L, M) Low somatic SNV and MMEJ deletion mutation burden in Ewing sarcoma tumors (ES; n = 23) as compared to neuroblastoma tumors (NB; n = 47). Short somatic mutations were determined from the whole genome sequencing data of tumors and matched normals. Each dot in K represents the total number of somatic SNVs for each tumor. Each dot in L represents the proportion of 3+-bp deletions with 2+-bp microhomology among all indels of each tumor.

To evaluate the impact of exon-skipping events on POL0 protein expression, we transduced sgRNAs targeting individual exons. Targeting *POLQ* exon 7, exon 21, or exon 25 by CRISPR-Cas9-mediated gene knockout induced profound protein loss of POL0 as well as a reduction of MMEJ activity (**Fig.2C, D**). Moreover, a cycloheximide chase assay revealed that a shorter isoform of the POL0 protein, expressed from a *POLQ* cDNA lacking exon 25, is more unstable than the full-length POL0 protein (**Fig. 2E**). Exon 25 skipping causes premature termination of POL0 protein synthesis, preventing the complete expression of the “Fingers” and “Palm” of the encoded polymerase. Structural modeling by AlphaFold 3 demonstrated that this truncation disrupts the structural integrity of this domain, likely explaining the instability compared to the full-length protein (**Fig. S3K**). The *POLQ* cDNA lacking exon 25 or exon7 failed to rescue the MMEJ-deficiencies in *POLQ* knockout cells (**Fig. 2F**).

To further investigate the biological significance of *POLQ* exon 25 skipping in the MMEJ activity, we designed antisense oligonucleotides (ASOs) that target exon 25 and induce exon 25 skipping (**Fig. S4A**). *POLQ* exon 25 skipping decreased POLΘ protein expression and MMEJ activity (**Fig. S4B, C**). Moreover, a mNeongreen-based minigene reporter, as previously described ^28^, was generated in which the mNeongreen is split into two exons with synthetic genes derived from the partial intron 24/25 and exon 25 of *POLQ* inserted in between (**Fig. S4D**). The minigene reporter recapitulated the *POLQ* exon 25 skipping driven by depletion of EWSR1, indicating a direct involvement of EWSR1 in the reduction of *POLQ* exon 25 skipping by binding the *POLQ* pre-mRNA near the exon 25 (**Fig. 2G, Fig. S4E**). Using an RNA immunoprecipitation protocol ^29^, we determined whether EWSR1 interacts with the *POLQ* pre-mRNA, using HEK293T cells harboring the FKBP12^F36V^-tagged EWSR1 (**Fig. S4F**). Indeed, EWSR1 efficiently co-immunoprecipitated with the *POLQ* pre-mRNA near exon 7 and exon 25, as compared to the *GAPDH* control (**Fig. S4G, H**). Taken together, EWSR1 binds directly to *POLQ* pre-mRNA and prevents exon skipping of the *POLQ* transcript, leading to maintenance of POL0 protein levels and MMEJ activity.

Given that one allele of *EWSR1* is lost in EwS cells, we speculated that these phenomena would also be observed in EwS cells, including patient samples. A higher level of exon 7 and exon 25 skipping and decreased expression of POLΘ protein were evident in EwS cell lines, compared to the osteosarcoma cell line U2OS (**Fig. 2H, I**). Importantly, patient-derived EwS tumor samples exhibited a higher frequency of *POLQ* exon 25 skipping than primary tumors from other pediatric solid tumors, including Neuroblastoma, Osteosarcoma, and Wilms tumor (**Fig. 2J**). Moreover, *POLQ* mRNA levels in Ewing sarcoma were comparable to or lower than other pediatric solid tumors (**Fig. 2K**).

We also evaluated single nucleotide variants (SNVs), the mutational signature, and small deletions in patient-derived EwS tumors. Consistent with previous studies ^30^, the EwS tumors have a low SNV frequency compared to Neuroblastoma, consistent with a MMEJ deficiency in the tumors (**Fig. 2L**). Although the level of HRD hallmarks, ID-6 and signature 3, were overall low, which is consistent with a previous study ^31^, Ewing sarcoma tumors showed an even lower level of signature 3 or ID-6, compared to Neuroblastomas, again confirming that EwS is an HR-proficient tumor **(Fig. S5A-F**). In contrast, the indels observed in the EwS samples, compared with neuroblastoma samples, have a lower frequency of deletions with >3bp micro-homologies and templated insertions, consistent with a low level of *POLQ*-mediated MMEJ in EwS ^32^ **(Fig. 2M, Fig. S5G-I**). To validate our findings in the primary patient samples, we measured the degree of microhomology at breakpoints of the *EWSR1* fusion in EwS tumor samples from two different EwS patient cohorts. Interestingly, the degree of microhomology at canonical translocation breakpoints was very low in the EwS genomes, consistent with a cellular defect in the MMEJ pathway and a compensatory increase in NHEJ (**Fig. S5J, K**). Collectively, EwS tumor cells have a bona fide defect in MMEJ, which is probably due to increased *POLQ* exon 25 skipping.

### The Prion-like domain of EWSR1 is required for the faithful splicing of the *POLQ* transcript

To identify the functional domain of EWSR1 required for accurate *POLQ* splicing, we next generated EWSR1 mutants lacking individual domains (**Fig. 3A**). Deletion of the prion-like domain (M1) or the C-terminus RRG2 domain (M6) resulted in failure to rescue the *POLQ* exon skipping and POL0 protein expression level (**Fig. 3B, C, Fig. S6A**). Consistent with these results, MMEJ activity was not restored in *EWSR1*-depleted cells by these two mutant forms of EWSR1 (**Fig. S6B**). In addition, these mutant proteins were unable to rescue the cellular hypersensitivity to topoisomerase inhibitors or an ATR inhibitor. The mutants also failed to rescue the synthetic lethality resulting from BRCA2 loss in *EWSR1*-deficient cells (**Fig. S6C-G**).

**Figure 3:**
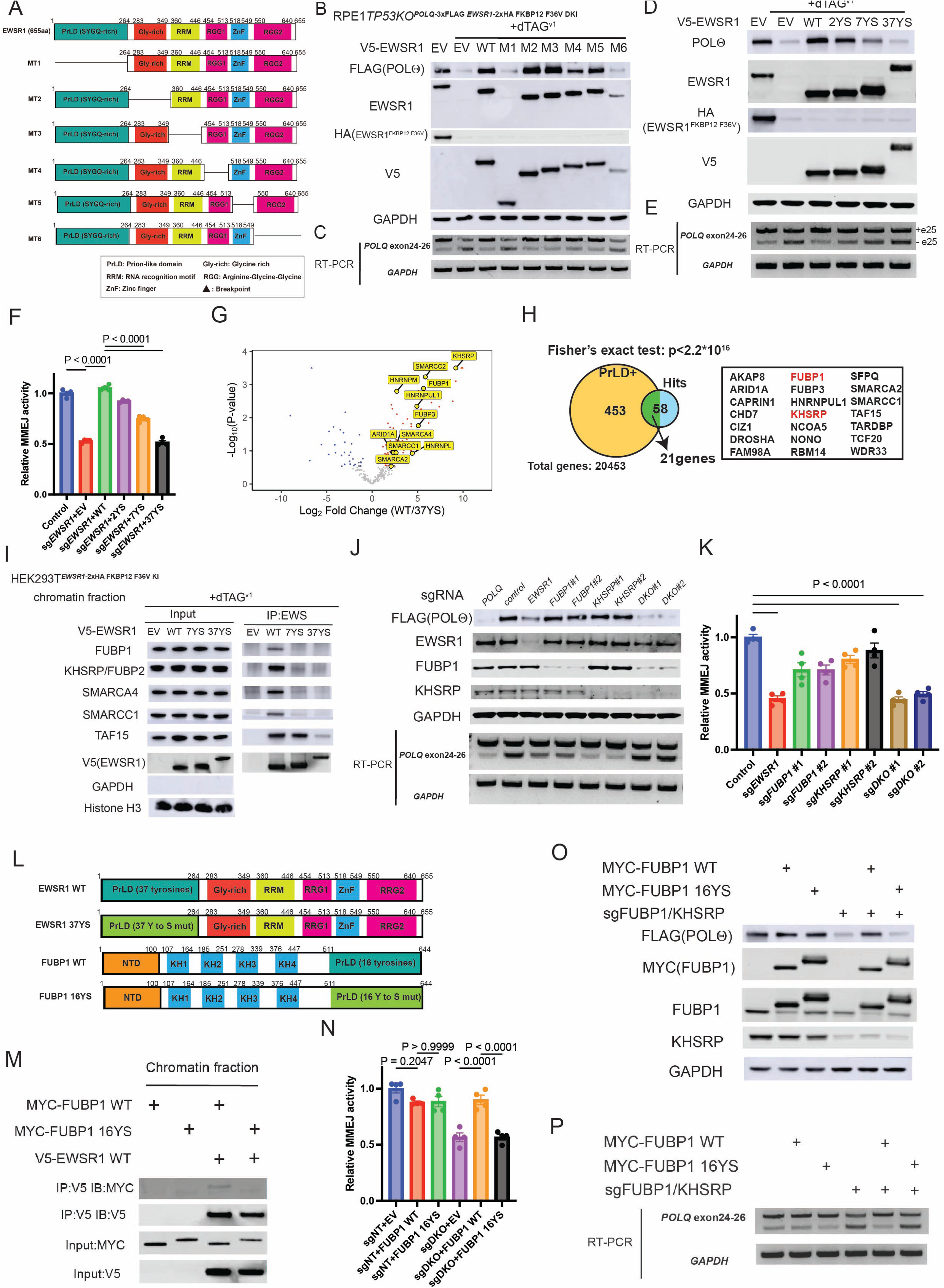
Formation of EWSR1 condensates through the prion-like domain is critical for faithful splicing of *POLQ* transcript via interacting with FUBP1 and KHSRP/FUBP2. **(A)** Schematic representation of the individual domains of EWSR1 and the deletion mutants used in Fig.3B, C. (B, D) RPE *p53*-/-3xFLAG-*POLQ*/2xHA-FKBP12 V36F *EWSR1* double knock-in cells were retrovirally transduced with empty vector, wildtype EWSR1, or indicated EWSR1 mutants. After transduction, cells were treated with blasticidin for four days. Then, cells were treated with DMSO or 1μM dTAG^v1^ for 24hr. After dTAG^v1^ treatment, cell lysates were subjected to immunoblot analysis. (C, E) RT-PCR showing the *POLQ* exon 25 skipping in the indicated cells. (F) U2OS MMEJ ver.2 (Cas9-based) cells were lentivirally transduced with control sgRNA or sgRNA targeting *EWSR1* together with empty vector wild-type EWSR1 or the indicated EWSR1 mutants by retroviral transduction. Cells were treated with puromycin and blasticidin for 4days, and then subjected to MMEJ reporter analysis (n=4). (G) Volcano plots comparing detected protein levels by IP-MS between EWSR1 wildtype and the EWSR1 37YS mutant. (H) A ven diagram showing the frequency of all the genes and positive hits with a prion-like domain identified by PLAAC. (I) Immunoprecipitation followed by immunoblotting analysis showing wildtype EWSR1 preferentially bound to FUBP1, KHSRP, SMARCA4 and SMARCC1 compared to the EWSR1 mutants. (J) FUBP1/KHSRP DKO cells exhibited POL0 protein reduction (upper) and increased *POLQ* exon 25 skipping (lower). (K) MMEJ activity was profoundly decreased in FUBP1/KHSRP DKO cells measured by U2OS MMEJ ver.2 cells (n=4). (L) Schematic representation of wildtype and YS-mutated EWSR1 or FUBP1. (M) Immunoblotting showing wildtype FUBP1 preferentially bound to EWSR1 compared to the FUBP1 16YS mutant. (N) U2OS MMEJ ver.2 cells were lentivirally and retrovirally transduced with indicated sgRNA(s) (co-expressing puromycin resistance gene) and control or FUBP1 cDNA (co-expressing blastidcidin resistance gene). Cells were treated with puromycin and blasticidin for 4days, and then subjected to MMEJ reporter analysis (n=4). (O) RPE *p53*-/-3xFLAG-*POLQ*/2xHA-FKBP12 V36F *EWSR1* double knock-in cells were lentivirally transduced with Cas9 and a sgRNA targeting control (NT) or sgRNAs targeting FUBP1 and KHSRP together with retrovirally transduced with a control vector, wildtype FUBP1 or the FUBP1 16YS mutant. After puromycin and blasticidin selection, cells were subjected to immunoblotting analysis. (P) RT-PCR showing the *POLQ* exon25 skipping in the indicated cells.

A nuclear localization signal (NLS) is located at the RRG2 domain, and its loss is likely to destabilize the EWSR1 protein. The NLS mutant protein failed to rescue *POLQ* exon skipping and failed to restore POLΘ protein expression. To further evaluate the prion-like domain, the domain was subdivided into five parts, and mutant proteins lacking the individual regions were generated (**Fig. S6H**). Interestingly, all mutant forms were able to rescue the POLΘ protein level, albeit with some minor differences, suggesting that the domains have functional redundancies (**Fig. S6I)**. Since the prion-like domain of EWSR1 is critical for the formation of condensates of EWSR1 ^33^, we infer that the formation of EWSR1 condensates may play a role in maintaining the faithful splicing of POL0. To test this hypothesis, we generated multiple mutant forms of EWSR1 in which tyrosine residues of the prion-like domain were substituted with serine residues, as previously described^33^. Strikingly, an EWSR1-7YS mutant, in which tyrosine residues were substituted by serine residues, failed to rescue the *POLQ* exon skipping and POLΘ protein expression level in *EWSR1*-depleted cells (**Fig. 3D, E, Fig. S6J, K**). Furthermore, an EWSR1-37YS mutant, in which all thirty-seven tyrosine residues in the prion-like domain were replaced with serine residues, and known to be defective in condensate formation, abolished the ability of EWSR1 to prevent exon skipping of the *POLQ* transcript, further suggesting that condensate formation is critical for faithful splicing. Consistent with these findings, the EWSR1-7YS and EWSR1-37YS mutants failed to rescue MMEJ activity in EWSR1-depleted cells (**Fig. 3F**). Taken together, the prion-like domain of EWSR1 is required for the faithful splicing of the *POLQ* transcript and for the expression of the POLΘ protein, suggesting that condensate formation of EWSR1 is also required for these activities as previously described ^5,33^.

### The prion-like domain of EWSR1 is critical for interaction with spliceosomes including FUBP1 and KHSRP

The condensate-defective EWSR1 mutant may have lost the ability to bind the *POLQ*-pre mRNA. However, RNA-immunoprecipitation revealed that the EWSR1-7YS and EWSR1-37YS mutants can still bind to the *POLQ* pre-mRNA (**Fig. S4 F-I**). Alternatively, the condensate-defective EWSR1 mutant may be competent for *POLQ* mRNA binding but defective in its interaction with RNA-binding proteins in spliceosomes. To test this hypothesis, we performed an IP/mass spectrometry analysis using HEK293T cells with a biallelic FKBP12-V36F knock-in at the N-terminus of *EWSR1* (**Fig. 3G**). The cDNAs encoding the control V5-tagged *EWSR1*-WT, or the *EWSR1* 37YS mutant, were transduced into the cells and, following dTAG^v1^ treatment, cellular proteins were immunoprecipitated with the anti-V5 antibody. A previous study demonstrated that the prion-like domain of EWS-FLI1 interacts with BAF complex subunit components in condensates ^5^. Accordingly, wildtype EWSR1, but not the EWSR1 37YS mutant protein, preferentially bound to BAF subunits, including ARID1A, SMARCA2, SMARCA4, and SMARCC1 (**Fig. 3G, I, Fig. S7A**). In addition to these BAF subunits, additional RNA binding proteins were identified as binding partners of wild-type EWSR1. Intriguingly, the interactors contain prion-like domains, further indicating a functional interaction of EWSR1 with these proteins in spliceosomes (**Fig. 3H**).

Since exon skipping in *EWSR1* KO cells is typically observed for exons that are flanked by longer introns (**Fig. S7B**), we focused on FUBP1, among the prion-like domain-dependent EWSR1 binding partners. FUBP1 is a spliceosome known to mediate the splicing of long introns ^34^ (**Fig. 3H, I**). In addition, KHSRP/FUBP2, known to have a similar function in splicing, was further evaluated. Depletion of FUBP1 or KHSRP alone cause only a minor reduction in POL0 protein expression or MMEJ activity. However, dual depletion of FUBP1 and KHSRP increased *POLQ* exon25 skipping, reduced the POL0 protein level, and reduced the cellular MMEJ activity, indicating a functional redundancy between FUBP1 and KHSRP (**Fig. 3 J, K, Fig. S7C).** A FUBP1-16YS mutant, with tyrosine residues of its C-terminus region substituted with serine residues, failed to bind to wildtype EWSR1 and failed to rescue the phenotype of FUBP1/KHSRP DKO cells, suggesting that tyrosine residues of EWSR1 and FUBP1 are critical for the protein-protein interaction and for *POLQ* exon25 inclusion (**Fig. 3 L-Q, Fig. S7D)**. Similarly, a KHSRP-17YS mutant failed to rescue the phenotype of FUBP1/KHSRP DKO cells **(Fig. S7E-H)**. Taken together, the protein-protein interaction between EWSR1 and FUBP1 or KHSRP via the prion-like domain is critical for the faithful splicing of the *POLQ* pre-mRNA.

### The EWS-FLI1 oncoprotein inhibits formation of POLQ mitotic foci at DSB loci and disrupts MMEJ

The chromosomal translocation of 11;22 leads not only to the loss of one wildtype EWSR1 allele but also to the generation of an oncogenic fusion protein EWS-FLI1 in EwS cells (**Fig. S8A**). Moreover, EWS-FLI1 is known to heterodimerize with EWSR1 and to disrupt its activity ^5,35,36^. We therefore hypothesized that this oncoprotein may directly or indirectly interfere with MMEJ activity. To test this hypothesis, the EWS-FLI1 oncoprotein was lentivirally expressed in U2OS-based MMEJ reporter cells and evaluated for MMEJ activity. Interestingly, EWS-FLI1 diminished the MMEJ activity in U2OS MMEJ reporter cells (**Fig.4A, Fig. S8B**). Consistent with these results, EwS cell lines exhibited resistance to POLΘ inhibitors, Novobiocin and ART558 ^37^, compared with osteosarcoma cells U2OS (**Fig. S8C, D)**. The influence of various mutant forms of the EWS-FLI1 fusion oncoprotein on MMEJ activity was tested (**Fig. 4B**). Expression of the SYGQ1-FLI1, SYGQ2-FLI1, ΔSYGQ1, or ΔSYGQ2 mutant by cDNA-mediated transfection decreased the MMEJ activity in the U2OS reporter cells, indicating functional redundancies in the prion-like domain of EWS-FLI1. Interestingly, the EWS-FLI1 37YS mutant, which failed to disrupt MMEJ activity (**Fig. 4B, Fig. S8B**), is defective in condensate formation and in binding to wild-type EWSR1 protein ^5,36^. Moreover, expression of EWS-FLI1, but not the EWS-FLI1 37YS mutant, conferred hypersensitivity to topoisomerase inhibitors (**Fig. S8E, F**).

**Figure 4:**
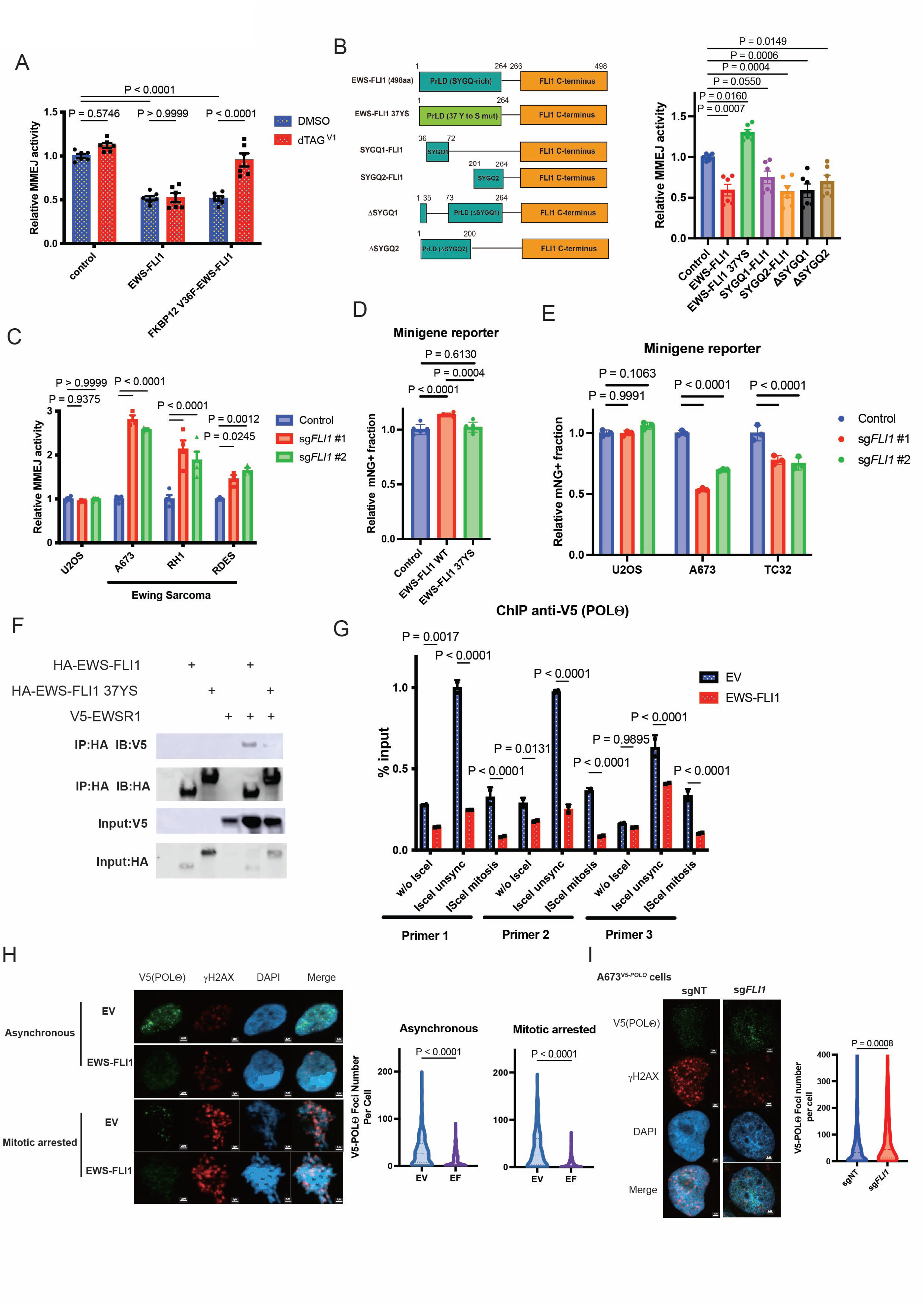
Expression of EWS-FLI1 perturbs POL0 foci assembly, resulting in impairment of MMEJ. (A) I-SceI-based MMEJ reporter cells were lentivirally transduced with an empty vector, wild-type *EWS-FLI1* or the FKBP12 V36F-tagged *EWS-FLI1*. Cells were treated with 1uM dTAG^V1^ for 18hr, and then adenovirally transduced with I-sceI. 48hr after I-sceI transduction, the frequencies of GFP positive cells were assessed by FACS (n=6). (B) Schematic representation of the EWS-FLI and mutant forms of EWS-FLI1 used in MMEJ reporter assay (left). I-SceI-based MMEJ reporter cells were lentivirally transduced with an empty vector, wild-type *EWS-FLI1* or indicated mutant forms of *EWS-FLI1* (co-expressing puromycin resistance gene). After puromycin selection, cells were subjected to MMEJ reporter assay (right, n=6). (C) I-SceI based MMEJ reporter was lentivirally transduced with U2OS, A673, RH1 and RD-ES cell lines. Cells were subsequently transduced with Cas9 together with control sgRNA or sgRNA targeting FLI1. Cells were treated with puromycin for 4days, and then subjected to MMEJ reporter analysis (n=3). (D) EWS-FLI1 but not the EWS-FLI1 37YS mutant increased *POLQ* exon 25 skipping measured by K562 minigene reporter cells (n=6). (E) A673 and TC32 cells carrying the minigene reporter showing depletion of EWS-FLI1 decreased *POLQ* exon 25 skipping (n=3). (F) Immunoprecipitation followed by immunoblotting analysis showing wildtype EWS-FLI11 preferentially bound to EWSR1 compared to the EWS-FLI1 37YS mutant. (G) ChIP-qPCR assay showing expression of EWS-FLI1 perturbed POL0 assembly to the DSB loci induced by Isce-I (n=2). (H) Immunofluorescence analysis showing endogenous V5-tagged POL0 foci formation was perturbed by EWS-FLI1 expression both asynchronous and mitotic arrested cells. Cells were treated with 2Gy irradiation and then incubated for 4hr (scale bars = 2μm). (I) Immunofluorescence analysis showing endogenous V5-tagged POL0 foci formation was restored by EWS-FLI1 depletion in V5-tagged-*POLQ* knock-in A673 cells. Cells were treated with 2Gy irradiation and then incubated for 4hr (scale bars = 2μm).

To determine the effect of the EWS-FLI1 oncoprotein on MMEJ activity in Ewing sarcoma cells, we transduced an Isce-I-based MMEJ reporter into the A673, RH1, and RD-ES EwS lines. As expected, depletion of EWS-FLI1 from these cells by sgFLI1, as confirmed by western blot, restored MMEJ activity (**Fig. 4C, Fig. S8G**). These EwS lines, with EWS-FLI1 depletion, exhibited resistance to the POLΘ inhibitor, ART558 (**Fig. S8H),** consistent with their restoration of MMEJ following EWS-FLI1 depletion. The minigene reporter assay showed that expression of EWS-FLI1 but not the EWS-FLI1 37YS mutant increased the *POLQ* exon 25 skipping, whereas depletion of EWS-FLI1 in EwS cells ameliorated the *POLQ* exon 25 skipping (**Fig. 4D, E)**. Consistent with a previous study ^5^, EWS-FLI1 preferentially bound to EWSR1 compared with the EWS-FLI1 37YS mutant, suggesting an inhibitory effect of EWS-FLI1 on EWSR1-mediated splicing (**Fig. 4F**). Given that R-loop accumulation driven by the EWS-FLI1 expression inhibits the recruitment of BRCA1 to the DSB loci, we asked whether R-loop accumulation may be involved in the MMEJ reduction in Ewing sarcoma. Expression of wildtype RNaseH1 or mutant RNaseH1 did not affect EWS-FLI1-mediated MMEJ reduction (**Fig. S8I, J**).

Recent studies indicate that DSB repair by POLΘ and MMEJ occurs during mitosis of the cell cycle ^38^. POL0 is phosphorylated by PLK1 and assembles with TOPBP1, along with Rhino and the 9-1-1 complex, into POL0 foci ^18,39^. To assess the influence of EWS-FLI1 expression on the assembly of DSB loci, we engineered endogenously V5-tagged POLΘ expressing MMEJ reporter U2OS cells by CRISPR/Cas9-mediated knock-in. After generation of adenoviral I-SceI-induced DSBs, we performed chromatin-immunoprecipitation (ChIP) with an anti-V5 antibody, followed by qPCR analysis. Overexpression of EWS-FLI1 in these cells disrupted the assembly of POL0 with sites of DNA double-strand breaks **(Fig. 4G).** Consistent with this result, MMEJ activity and the assembly of POLΘ foci assembly, as measured by immunofluorescence, was decreased by the overexpression of EWS-FLI1 (**Fig. 4H**). Moreover, EWS-FLI1 overexpression in these U2OS cells also increased γH2AX foci, consistent with the loss of MMEJ activity in mitosis. EWS-FLI1 overexpression can therefore reduce POLΘ-mediated DSB repair in mitosis, at a time when POLΘ-mediated MMEJ is highly active ^18,39^. EWS-FLI1 was also depleted in endogenously V5-tagged POLΘ expressing A673 cells. As predicted, the loss of EWS-FLI1 restored POL0 foci assembly (**Fig. 4I).** Collectively, the EWS-FLI1 oncoprotein can reduce the level of POLΘ mitotic foci, leading to a functional impairment of MMEJ in EwS cells.

### Inhibition of the FA, NHEJ, or HR pathways efficiently kills the MMEJ-deficient Ewing sarcoma cells via synthetic lethality

Recent studies have shown that targeting the Fanconi Anemia (FA) pathway is synthetic lethal with MMEJ deficiency^40–43^. We therefore hypothesized that MMEJ-defective Ewing sarcoma cells would exhibit high dependence on the FA/BRCA signaling pathway. Indeed, depletion of FANCA, FANCI or FANCL efficiently reduced colony forming activity in the control A673 cells, whereas it showed only a minor impact on the EWS-FLI1-depleted A673 cells (**Fig. 5A**). Moreover, depletion of the EWS-FLI1 oncoprotein reduced FANCD2 foci assembly with Mitomycin C (MMC) treatment in Ewing sarcoma cells, A673 and TC32 cells, whereas sgFLI1 did not affect the activation of the pathway in U2OS cells (**Fig. 5B, S5A, B**). Since RBM39 degraders are potent FA pathway inhibitors, by inducing intron retention of the *FANCD2* pre-mRNA^44^, we tested the effect of RBM39 degraders, E7820 and Indisulam, on the FA pathway in Ewing sarcoma cells, A673 cells. Consistently, RBM39 degraders decreased FANCA and FANCD2 protein expression in A673 cells (**Fig. 5C**). Intriguingly, RBM39 degraders eliminated Ewing sarcoma cells efficiently (**Fig. 5D, E**), which can be rescued by depletion of the EWS-FLI1 oncoprotein in A673 and TC32 cells (**Fig. 5F, G, S5C, D**). These data indicate that targeting the FA/BRCA pathway by the RBM39 degraders is a promising therapeutic strategy for Ewing sarcoma cells.

**Figure 5:**
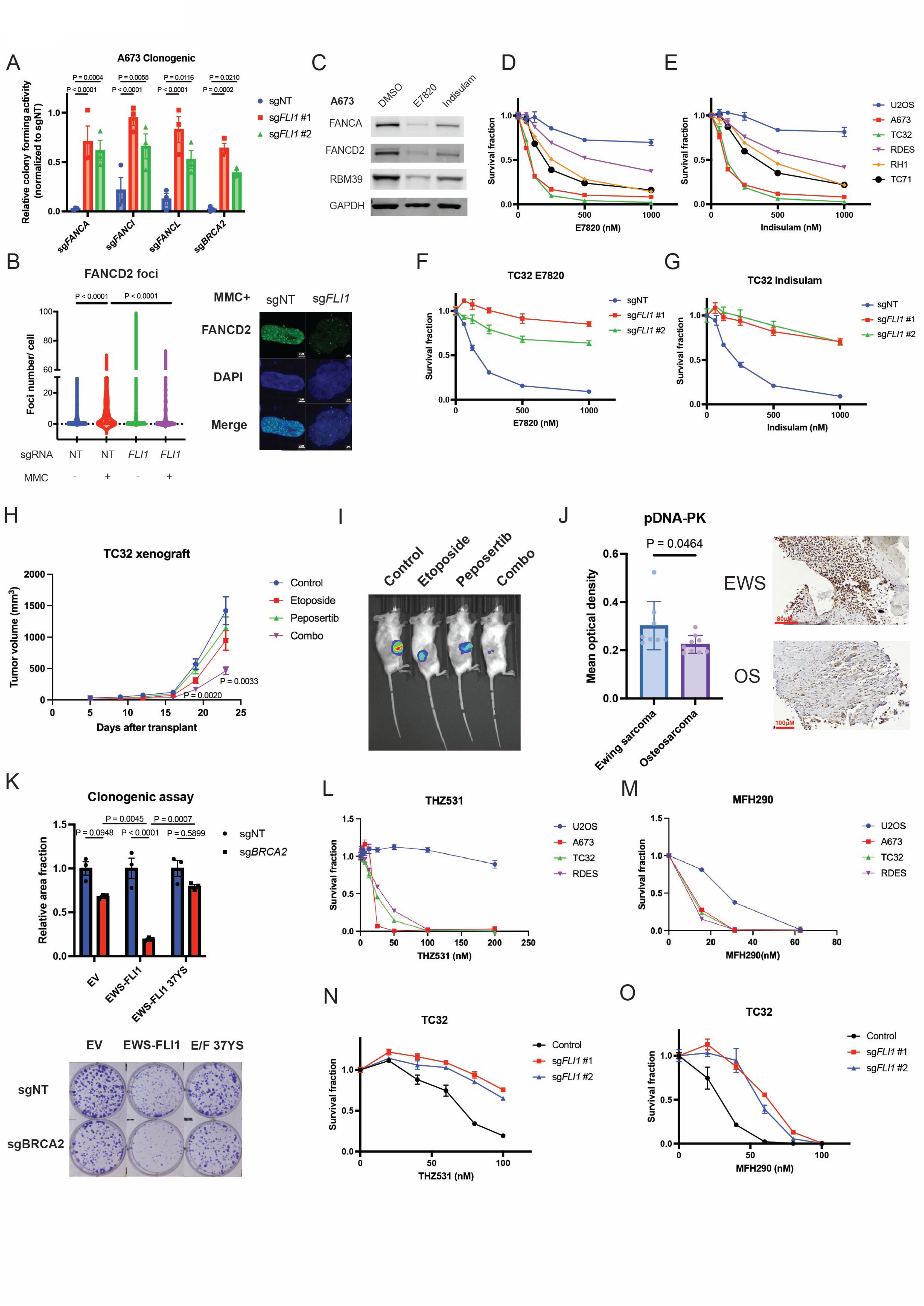
Inhibition of FA, NHEJ or HR efficiently eliminates Ewing sarcoma cells. (A) Colony forming assay showing that the control A673 cells exhibited higher dependence on *FANCA*, *FANCI*, *FANCL*, and *BRCA2* genes compared with the EWS-FLI1-depleted A673 cells by sg*FLI*1. (B) Immunofluorescence analysis showing FANCD2 foci formation was increased by EWS-FLI1 expression in A673 cells. Cells were treated with 10ng/ml MMC for 24hr. Shown are the number of FANCD2 foci (left) and the representative images of the cells treated with MMC (right, scale bars = 2um). (C) Immunoblotting analysis of the cell lysates from A673 cells treated with DMSO, 250nM E7820 or 250nM Indisulam for 72hr. Treatment with RBM39 degraders decreased FANCA and FANCD2 protein expression. (D, E) The CTG assay showing that compared with U2OS, Ewing sarcoma cells showed hypersensitivity to RBM39 degraders, E7820 (D) and Indisulam (E). (F, G) The CTG assay showing that depletion of EWS-FLI1 by sgFLI1 in TC32 cells led to acquisition of resistance to RBM39 degraders, E7820 (F) and Indisulam (G). (H, I) Xenograft model using TC32 showing therapeutic efficacy of a Peposertib and Etoposide. NSG mice (n = 10 mice/group) bearing luciferized TC32 cells were treated with 100 mg/kg of Peposertib (orally), 8 mg/kg etoposide (via IP), or both for 2 weeks. Tumor sizes were measured every 2-3 days (H) and also monitored by bioluminescence images (I). (J) Immunohistochemistry analysis from the patient samples of EwS (n=8) showing higher pDNA-PKc activities compared with that of OS (n=9). (K) Colony formation assay data of RPE *p53*-/-cells transduced with a control vector, EWS-FLI1 or EWS-FLI1 37YS together with or without BRCA2 depletion (n=3, top) and the representative colony images (bottom). (L, M) The CTG assay showing that compared with U2OS, Ewing sarcoma cells showed hypersensitivity to CDK12 inhibitors, THZ531 (L) and MFH290 (M). (N, O) The CTG assay showing that depletion of EWS-FLI1 by sg*FLI*1 in TC32 cells led to acquisition of resistance to CDK12 inhibitors, THZ531 (N) and MFH290 (O).

Since inhibiting two of three DSB repair pathways is synthetically lethal in cells ^45,46^, we asked whether inhibition of NHEJ or HR would also provide a viable therapeutic modality in MMEJ-deficient Ewing sarcoma cells. Previous studies have shown that inhibitors of NHEJ are synthetic lethal with MMEJ loss ^45,46^. Consistent with our study, EwS cells are sensitive to DNA-PK inhibitors, known to inhibit NHEJ and enhance the activity of cytotoxic agents in these cells^47^. We confirmed that EwS cell lines are sensitive to the DNA-PKci (**Fig. S5E**) and have an elevated baseline level of activated pDNA-PKc (**Fig. S5F**). EwS cells exhibited high phospho-RPA, a replication stress biomarker, and were hypersensitive to the topoisomerase inhibitors, topotecan, and etoposide (**Fig. S8A, S9G, H**). Since etoposide is used in dose-intensive chemotherapy of Ewing sarcoma, we asked whether inhibition of NHEJ could increase the therapeutic efficacy of etoposide in EWS cells. Strikingly, the DNA-PK inhibitor, Peposertib, enhanced the activity of etoposide in two EWS cell lines (**Fig. S9I, J)**. Moreover, the EwS cell line A673, grown as a xenograft, was also sensitive to peposertib, *in vivo*, and the drug enhanced the activity of etoposide (**Fig. 5H, I, Fig. S9K, L**). Consistent with the enhanced sensitivity to peposertib, immunohistochemistry analysis of the EwS primary patient samples revealed an elevated baseline level of pDNA-PKc compared with Osteosarcoma patient samples (**Fig. 5J**).

On the other hand, EWS-FLI1-but not EWS-FLI1-YS mutant-expressing cells are more dependent on BRCA2, confirming that the HR pathway is also a rational target in MMEJ-deficient EwS cells (**Fig. 5A, K**). Previous studies have also suggested that EwS cells are sensitive to HR inhibitors, perhaps through a mechanism of synthetic lethality. CDK12 is a known activator of HR repair, based on its ability to promote the transcription of long HR genes^48^. A targeted inhibitor of CDK12 is known to enhance the killing of EwS cells ^49^, though the mechanism of this cell killing was unclear. Consistent with this report, EwS cell lines are sensitive to the CDK12 inhibitors THZ531 and MFH290 (**Fig 5 L, M**). Depletion of the EWS-FLI1 oncoprotein in the EwS cells resulted in a restoration of MMEJ activity and acquisition of CDK12i resistance (**Fig 5N, O, Fig. S9M, N**). Taken together, the killing of the EwS cells by an HR inhibitor results mechanistically from synthetic lethality between the HR pathway and the MMEJ pathway.

## DISCUSSION

Despite the high sensitivity to DNA-damaging agents, EwS cells, expressing the EWS-FLI1 fusion oncoprotein, have a remarkably low somatic mutation burden ^30,50,51^. This observation suggested that EwS cells are deficient in an error-prone DNA repair mechanism, such as MMEJ. Indeed, CRISPR screens confirmed that the knockout of the EWSR1 gene functionally resembles the knockout of known MMEJ genes, such as *POLQ*, *XRCC1*, *APEX2*, and *RHNO1*, suggesting that EWSR1 protein cooperates with these other encoded proteins in the MMEJ pathway. Moreover, EWSR1-deficient cancer cell lines and EwS tumors have a low ID6 mutational signature, indicating a low level of microhomology-mediated repair of double-strand breaks. Since EWSR1 has a known function as a mRNA splicing factor ^35,52–54^, we hypothesized that EWSR1 may regulate MMEJ indirectly through its splicing of the mRNA encoding a MMEJ protein. EWSR1 is indeed required for the accurate splicing of the long mRNA encoding the critical MMEJ polymerase, POLΘ, although it is not required for splicing of mRNAs encoding other MMEJ proteins. Disruption of EWSR1 activity in EwS cells results in skipping of multiple exons in the *POLQ* pre-mRNA, especially exon 25, leading to decreased POLΘ expression in EwS cells and to a strong MMEJ defect.

In contrast to previous reports, our study indicates that EwS tumors are competent for high-fidelity DNA repair pathways, such as the homologous recombination pathway, and the tumors are PARP inhibitor-resistant. Other recent studies indicate that EwS cells are dependent on the HR genes, BRCA1, BARD1, and PALB2 ^55,56^ for survival. Previous studies have also demonstrated that HR deficiency is synthetic lethal in cells with a MMEJ deficiency ^22,41,57^. Accordingly, the strong MMEJ defect in EwS cells could result in an increased reliance on HR repair.

The chromosome 11/22 translocation in EwS cells and the generation of the EWS-FLI1 fusion oncoprotein appears to account, at least in part, for the cellular defect in EWSR1 splicing activity. First, since one allele of EWSR1 is disrupted by the chromosome translocation, the EwS cells already have a 50% reduction in EWSR1 protein expression. Second, the EWS-FLI1 oncoprotein heterodimerizes with EWSR1 and exerts a dominant negative effect on EWSR1 function ^5^. Our studies indicate that loss of EWSR1 function can account for the decreased expression of POL0 protein, the elevated exon 25 skipping of the *POLQ* pre-mRNA, and the defective MMEJ activity in EwS cells. Enhanced expression of the EWSR1 protein, or acute knockdown of the EWS-FLI1 oncoprotein, can reverse these defects and restore MMEJ activity, thereby reducing the cellular reliance on other DSB repair pathways, such as HR and NHEJ.

Splicing of specific mRNAs is regulated, at least in part, by RNA Binding Proteins (RBPs) which function in membrane-less condensates. Condensates known as nuclear speckles are required for splicing activity^58^. Many RNPs, including the EWSR1 protein, contain a prion-like domain (PrLD), allowing the proteins to assemble with other RBPs contained in condensates. We reasoned that EWSR1 might interact with other RBPs and function cooperatively as a spliceosome for the *POLQ* pre-mRNA. Moreover, disruption of the PrLD of EWSR1, through the generation of multiple Tyrosine to Serine (YS) mutations, disrupts the assembly of functional spliceosomes. Indeed, the EWSR 37YS mutant, known to be defective in condensate assembly ^5^, also fails to correct exon 25 skipping of the *POLQ* mRNA, and fails to restore POL0 protein expression. IP/Mass Spectrometry identified specific RBPs that interact with EWSR1. Interestingly, multiple RBPs containing prion-like domains were identified in the immune complexes, and these proteins did not coimmunoprecipitate with the EWSR1 37YS mutant protein. Components of SWI/SNF complexes, including subunits of the cBAF complex with prion-like domains, were also found to interact with EWSR1 and EWS-FLI1, consistent with a previous report ^5^. Unlike EWSR1, other FET proteins containing PrLD domains, such as FUS and TAF15, were not required for accurate *POLQ* splicing, suggesting that EWSR1 forms distinct RBP complexes, perhaps functioning in distinct condensates. Taken together, EWSR1 may act as a “hub” for spliceosomes, using its specific prion-like domain for binding to a distinct set of RBPs and for splicing of the *POLQ* mRNA and probably other mRNAs. Expression of EWS-FLI1 disrupts this splicing mechanism.

Some RNA binding proteins, including FUBP1 and KHSRP/FUBP2, specifically interact with wildtype EWSR1, but not with the EWSR1 37YS mutant. FUBP1 and KHSRP have high similarity in their protein sequence and exhibit functional redundancy in their splicing activity ^59^. FUBP1 and KHSRP are multifunctional RNA binding proteins and are known to redundantly regulate the splicing of the SMN2 mRNA ^59^. A carboxy-terminal, tyrosine-rich region is present in both FUBP1 and KHSRP/FUBP2 ^60^. However, the biological functions of the tyrosine-rich regions of FUBP1 and KHSRP have remained unclear. Since the tyrosine residues of the prion-like domain are critical for liquid-liquid phase separation^61,62^ and condensate formation, a YS mutant of FUBP1 or KHSRP, may fail to form condensates with EWSR1 and may lead to aberrant splicing events, such as *POLQ* exon skipping. FUBP1 is crucial for splicing long introns ^34^ such as the long intron 24/25 in the *POLQ* pre-mRNA. Therefore, a condensate formed by EWSR1-FUBP1/KHSRP may be required for the accurate splicing of exons trapped between long introns. Interestingly, the dual knockout of FUBP1 and KHSRP, like the knockout of EWSR1, results in exon 25 skipping of the *POLQ* mRNA and a reduction in POL0 protein expression and MMEJ. Re-expression of the wildtype FUBP1, but not of the FUBP1-YS mutant, rescued these defects in the dual knockout cells. The RBPs, EWSR1, FUBP1, and KHSRP therefore appear to functionally interact via their PrLD domains and to cooperate in a EWSR1-initiated splicing pathway for the *POLQ* mRNA (**Fig. S10**).

Given that overexpression of EWS-FLI1 oncoprotein induces extensive DNA damage^63^, we reasoned that the EWS-FLI1 fusion oncoprotein could impair MMEJ activity in EwS cells. Indeed, the acute degradation of the EWS-FLI1 oncoprotein rapidly rescued MMEJ, suggesting an acute upregulation of POL0 expression and POL0 mitotic foci. While the EWS-FLI1 protein forms condensate with EWSR1, it disrupts the function of EWSR1, reduces POL0 expression, and inhibits MMEJ. The EWS-FLI1-37YS mutant fails to form condensates with wild-type EWSR and fails to reduce MMEJ. While knockout of EWS-FLI1 restores MMEJ, loss of the oncoprotein is also detrimental to EwS cell growth and results in a reduction in the transcription of EWS-FLI1 target genes which contributes to EwS cell growth.

Previous studies on the pathogenesis of Ewing Sarcoma have focused primarily on the transcriptional activity of the EWS-FLI1 oncoprotein, driven by the carboxy-terminal FLI1 transcriptional domain. Indeed, this oncoprotein upregulates the transcription of genes which in turn promote R-loops and the neoplastic growth of the EwS tumor. Our study also provides new insights into the pathogenesis of the cancer. While the chromosome 11/22 translocation results in the formation of the EWS-FLI1 oncoprotein, it also simultaneously inactivates the EWSR1 *tumor suppressor* function. One allele of this EWSR1 tumor suppressor gene is inactivated by the translocation itself, and the remaining EWSR1 suppressor function is inactivated by the binding of the fusion oncoprotein to the remaining EWSR1 protein. Due to the loss of the splicing activity of EWSR1, the tumor cells exhibit the characteristic MMEJ defect-namely, radiation sensitivity, a defect in microhomology-mediated DSB repair, and a hypo-mutable phenotype. The splicing defect may also result in the inactivation of other tumor suppressor genes and a more direct role in the process of oncogenesis. For instance, the EWS-FLI1 fusion protein may have additional direct oncogenic functions beyond its ability to promote transcription, R-loops, and replication stress. Specifically, the EWS-FLI1 fusion can disrupt PLK1-activated POLΘ mitotic foci ^39^ which are known sites of TOPBP1-mediated DSB repair. Given that inhibition of MMEJ under replication stress leads to sensitization to PARPi and ATRi^64^, an MMEJ defect in Ewing sarcoma, despite its HR proficiency, may be responsible for its sensitivity to these drugs. Accordingly, the EWS-FLI1 oncoprotein may further block the repair of DSBs in mitosis, promote the formation of micronuclei, and enhance genomic instability.

Taken together, *Ewing sarcoma is the first known cancer with loss of MMEJ activity.* The impairment of the MMEJ pathway in EwS prompted us to target the FA, NHEJ, or HR pathways, the three DNA repair pathways known to be synthetic lethal with MMEJ loss. Indeed, EwS cells are sensitive to the RBM39 degraders, which lead to the disruption of the FA pathway. In addition, EwS cells are sensitive to a NHEJ inhibitor, peposertib, as a monotherapy. Peposertib is also synergistic with the genotoxic drug, Etoposide, in killing EwS cells. Accordingly, the use of this drug combination could reduce the need for high doses of Etoposide for EWS patients and thereby reduce its long-term toxicity, such as therapy-related leukemia^65^. EWS cells, due to their MMEJ defect, are also sensitive to the HR inhibitor, THZ531, a CDK12 inhibitor. This inhibitor is one of the first known inhibitors of the HR pathway. Exon 25 skipping of the *POLQ* mRNA in fresh tumor samples may provide a biomarker for identifying tumors with an underlying MMEJ defect which may in turn respond to targeted NHEJ inhibitors or HR inhibitors through the mechanism of synthetic lethality.

## EXPERIMENTAL MODEL AND SUBJECT DETAILS

### Cell cultures

RPE *p53*-/-cells generated previously^66^ were grown in DMEM/F12 containing GlutaMAX (Gibco) and supplemented with 10% fetal bovine serum (FBS) (Sigma-Aldrich). HEK293T (ATCC, CRL-3216) cells were grown in DMEM (Gibco) supplemented with 10%FBS. TC32 and TC71 cell lines were obtained from children cancer repository and grown in RPMI (Gibco) supplemented with 10%FBS. RH-1 (DSMZ, ACC493) cell grown in RPMI 1640 supplemented with 10%FBS, 1% P/S. RD-ES (ATCC, HTB-166) cells were grown in RPMI 1640 supplemented with 20%FBS, 1% P/S. SK-ES1 (ATCC, HTB-86) and SK-PN-DW (ATCC, CRL-2139) cells were cultured in DMEM (Gibco) supplemented with 10%FBS. U2OS parental (ATCC, HTB-96) and carrying a DNA repair template cells were grown in DMEM (Gibco) supplemented with 10%FBS. The BRCA1-deficient cell lines HCC1937 (ATCC, CRL-2336) and HCC1954 (ATCC, CRL-2338) and BRCA1-complemented cells were grown in RPMI (Gibco) supplemented with 10%FBS. RH-1 (DSMZ, ACC493). K562 (ATCC, CCL-243) cells were grown in RPMI 1640 supplemented with 20%FBS, 1% P/S. MDA-MB-436 and MDA-MB-436+BRCA1 cells were a kind gift from Dr. Geoffrey Shapiro (Dana Farber Cancer Institute) and grown in RPMI (GIBCO) supplemented with 10% fetal bovine serum. DLD1 BRCA2 -/-and DLD1 Parental were obtained from Horizon discovery, Waterbeach, UK. U2OS cells with HR reporter (DR-GFP) was a kind gift from Dr. Jeremy Stark (City of Hope).

To establish the U2OS-MMEJ reporter cells, U2OS cells were lentivirally transduced with the Isce-I-based (ver.1) or Cas9-based (ver.2) MMEJ reporter and then incubated with puromycin. After selection with puromycin, a single clone was isolated and subsequently the clones were validated. All cells were cultured in an incubator maintained at 37°C, 5% CO_2_ and a relative humidity of 95%.

### Compounds and antibodies

Chemical compounds, including Olaparib (#S1060), Novobiocin (#S2492), Etoposide (#S1225), VE822 (#S7102), Ro-3306 (#S7747), E7820 (#S6628), Indisulam (#S9742), M3814 (#S8586) and THZ0531 (#$6595) were purchased from Selleckchem. ART558 (#HY-141520) and MFH-290 (#HY-153244) were purchased from MedChemExpress. Nocodazole (#1404) was purchased from Sigma Aldrich. Chemicals were dissolved in DMSO and kept in small aliquots at −20°C or 80°C. The key sources of antibodies are: POLθ (Cell Signaling Technology, #64708), γH2AX (Millipore #05–636), LIG3 (BD Biosciences, #611876), HMCES (Santa Cruz Biotechnology, SC-514238), HMCES (Atlas Antibodies #HPA044968), APEX2 (Cell Signaling Technology, #74728), PARP1(Cell Signaling Technology, #9542), CIP2A (Abcam, ab128179), DNA-PK (Cell Signaling Technology, #12311), Phospho-DNA-PKcs (Ser2056) (Cell Signaling Technology, #68716), TOPBP1 (Abcam, ab2402), XRCC1 (Cell Signaling Technology, #76998), FEN1 (Cell Signaling Technology, #82354), EWSR1 (Cell Signaling Technology, #11910 for WB), EWSR1 (Santa Cruz Biotechnology, sc-28327 for WB), EWSR1 (Life Technologies, A300417A for IP and RIP), BRCA1 (Merck Millipore, OP92), BRCA2 (Cell Signaling Technology, #10741), FLI1 (Abcam, ab133485), FUS (Cell Signaling Technology, #67840), TAF15(Cell Signaling Technology, #28409), Vinculin (Cell Signaling Technology, #4650), FLAG (Sigma, F1804), HA (Cell Signaling Technology, #3724 for WB), HA (Roche, #11867431001 for IP and RIP), V5 (Cell Signaling Technology, #13202 for WB and IF), V5(Abcam, ab15828 for ChIP), V5(Abcam, ab27671 for IP and WB), GAPDH (Cell Signaling Technology, #5174), Histone H3 (Cell Signaling Technology, #9715), FUBP1 (Abcam ab181111), KHSRP (Cell Signaling Technology, #13398), SMARCA4/BRG1 (Cell Signaling Technology, #3508), SMARCC1 (Cell Signaling Technology, #11956), MYC (Santa Cruz Biotechnology, SC-40), FANCA (Cell Signaling Technology, #14657), FANCD2 (Novus Biologicals, NB100-182).

### Generation of CRISPR engineered cell lines

For generating *EWSR1* knock-out clones in HEK293T and RPE *p53*-/-cells, a ribonucleoprotein (RNP) complex was formed by Alt-R™ S.p. HiFi Cas9 and CRISPR-Cas9 sgRNA targeting *EWSR1* (Integrated DNA Technologies). For generating endogenously 3xFLAG-tagged *POLQ* knock-in HEK293T and RPE *p53*-/-cells, 3xFLAG-tagged *POLQ* HEK293T *EWSR1*-FKBP12 V36F double knock-in cells or V5-tagged *POLQ* knock-in U2OS and A673 cells, the Cas9/sgRNA RNP complex together with a template was delivered. Cells were resuspended in SE (U2OS cells), SF (HEK293T and A673 cells) or P3 (RPE *p53*-/-cells) Nucleofector Solution with supplement (Lonza) and then mixed with the Cas9/sgRNA RNP complex and Alt-R™ Cas9 Electroporation Enhancer (Integrated DNA Technologies), respectively. The Cas9/sgRNA RNP complex was delivered to the cells by 4D-Nucleofector^TM^ (Lonza). Media was changed the following day, and then 2 days after cells were seeded in a 96-well plate to isolate single clones. Individual clones were tested for genomic editing analyses using immunoblotting and genomic PCR with subsequent Sanger sequencing.

### Plasmids

psPAX2 and pMD2.G (Addgene #12260 and #12259) were gifts from Dr. Didier Trono. pLV-EF1a-IRES-Blast (pLV-Blast) (Addgene #85133) was a gift from Dr. Tobias Meyer.^67^ pCDH-puro-EWS-FLI1 was a gift from Jialiang Wang (Addgene plasmid # 102813). pLenti CMV Puro LUC (w168-1) was a gift from Eric Campeau & Paul Kaufman (Addgene plasmid # 17477). pCVL.SFFV.d14GFP.EF1a.HA.NLS.Sce(opt).T2A.TagBFP was a gift from Andrew Scharenberg (Addgene plasmid # 32627). EWS-FLI1 37YS mutant was cloned into pCDH-puro vector. EWSR1, FUBP1, KHSRP, RNaseH1 and its mutant forms were cloned into pMYs-IRES-Blastcidin vector, respectively. sgRNAs were cloned into a lentiCRISPR-v2 construct as previously described^68^. Of note, sgEWSR1 #2, sgFUBP1 #2 or sgKHSRP #2 target genomic DNA but not cDNA, so that endogenous knockout and ectopic cDNA expression are possible simultaneously. All the sgRNA sequences used in this study were provided in Supplemental Table 1.

### Virus generation

Lentivirus was generated by co-transfecting lentiviral constructs with packaging vectors, psPAX2 and pMD2.G, into HEK293T cells using CalPhos Mammalian Transfection Kit (Takara Bio). Two days after transfection, supernatant containing virus was harvested and filtered through 0.45 µm filter. Viral transductions were performed in the presence of 8 µg/ml (for RPE *p53*-/-, MDA-MB 436, DLD1, HCC1937, and HCC1954 cells) or 4 µg/ml (for K562 cells) polybrene (Sigma-Aldrich). For all virus generations and transductions media supplemented with heat inactivated FBS was used. Retrovirus was generated by co-transfecting retroviral constructs with packaging vectors, and, into HEK293T cells using CalPhos Mammalian Transfection Kit (Takara Bio). Two days after transfection, supernatant containing virus was harvested and filtered through 0.45 µm filter. Viral transductions were performed in the presence of 8 µg/mL polybrene (Sigma-Aldrich). Puromycin dihydrochloride (Sigma-Aldrich) or Blasticidin S HCl (Thermo Fisher Scientific) was used after infection of retrovirus or lentivirus.

### Colony formation assay

Cells were seeded in 6-well plates at 500-10,000 cells/well. The day after seeding, cells were exposed to each drug or x-ray. X-ray irradiation was performed using RS-2000 Irradiator (Rad Source Technologies). After incubation, cells were washed with PBS and then fixed in a mixture of 50% methanol, 10% acetic acid, and 40% water for 20 minutes. The fixed cells were stained with 0.5% crystal violet (Sigma-Aldrich) in 20% methanol for 2–6 h and then washed with water twice. The plates were imaged on GE Amersham Imager 600 using colorimetric transillumination setting. The area fraction of the colonies in each well was estimated using ImageJ.

### CellTiter-Glo assay

To test drug sensitivity, U2OS or Ewing sarcoma cells were seeded in 96-well plates at 1000–3000 cells/well. The day after seeding, drug was added to the wells, and then the plates were incubated for 3-5 days. After incubation, ATP in each well was quantified using CellTiter-Glo Luminescent Cell Viability Assay (Promega). CellTiter-Glo assays were performed in technical duplicate if not otherwise specified in a figure legend.

### Immunofluorescence

Mitotic synchronization was achieved using Ro-3306 (8 μM, 16 hours) followed by 30min release in fresh medium. Then cells were incubated with 100ng/ml nocodazole and subsequently irradiated. 4h after irradiation or in the absence of treatment, cells were fixed with 4% paraformaldehyde for 15 minutes at room temperature. For FANCD2 foci evaluation, cells were treated with 10ng/ml MMC for 24hr. Cells were then permeabilized with 0.3% Triton X-100 for 10 minutes on ice, followed by blocking with 3% non-fat milk for 1 hour at room temperature. The slides were stained with primary antibodies at 4℃ overnight. Afterward, they were stained with secondary antibodies for 1 hour at room temperature. The slides were scanned using fluorescence microscope. At least 100 cells were counted for each sample. Foci quantification was performed using software CellProfiler version 4.2.6.

### Immunoblotting

Whole cell lysate was prepared by lysing cells in ice-cold RIPA buffer (Cell Signaling Technology) supplemented with Protease/Phosphatase inhibitor Cocktail (Cell Signaling Technology). Cell lysate was cleared by centrifuging the samples at 12000 rpm, 4°C for 20 minutes. Supernatant was recovered and protein concentration was estimated using BCA assay kit (Thermo Fisher Scientific). Protein samples were boiled in 2X Laemmli sample buffer (Bio-rad) supplemented with 5% 2-mercaptoethanol or NuPAGE LDS sample buffer (Thermo Fisher Scientific) supplemented with NuPAGE sample reducing agent (Thermo Fisher Scientific). Samples were run in NuPAGE 4-12% Bis-Tris gels (Thermo Fisher Scientific) using NuPAGE MES SDS Running Buffer (Thermo Fisher Scientific) or NuPAGE 3-8% Tris-Acetate Gel (Thermo Fisher Scientific) using NuPAGE Tris-Acetate SDS Running Buffer (Thermo Fisher Scientific). After electrophoresis, samples were transferred onto a nitrocellulose membrane (Amersham). Membranes were blocked for 1 h with Blocking Buffer, which was prepared by diluting Fish Serum Blocking Buffer (Thermo Fisher Scientific) with Tris-buffered saline containing 0.1% Tween20 (TBST). Membranes were incubated with primary antibodies diluted in Blocking Buffer overnight at 4°C. Membranes were washed with TBST three times, 5 minutes each, incubated with secondary antibodies diluted in blocking buffer for 1h, which either horseradish peroxidase (HRP)-conjugated for chemiluminescent detection or fluorophore-conjugated for visualization using the LI-COR imaging system. After secondary antibody incubation, membranes were washed with TBST and imaged using chemiluminescence setting on GE Amersham Imager 600 or the LI-COR imaging system.

### DNA repair template assay

DR-GFP and EJ5-GFP cells were kind gifts from Dr. Jeremy Stark. MMEJ-GFP cells (ver.1 and ver.2) were generated by lentivirally transducing a cassette into U2OS or Ewing sarcoma cells using a pLV-EF1a-IRES-puro vector (addgene #85132) as previously described ^20^. U2OS cells carrying a DNA repair template reporter (DR-GFP, EJ5-GFP, and MMEJ-GFP) or Ewing sarcoma cells carrying a MMEJ reporter were lentivirally transduced with Cas9 and sgRNA (co-expressing puromycin resistance gene). After puromycin or blasticidin selection, 40,000 DNA repair template reporter cells were seeded in 12-well plates and adenovirally (for U2OS) or lentivirally (co-expressing BFP, for Ewing sarcoma cells) transduced with Isce-I the following day. 48h after Isce-I transduction, GFP signals were analyzed by CytoFLEX (Beckman). The signals were normalized to control cells. For U2OS Cas9-based MMEJ reporter assay, the cells were subsequently transduced with Cas9 and sgRNA targeting the reporter (co expressing BFP). 72h after Cas9/sgRNA transduction, GFP signals within BFP positive fraction were analyzed by CytoFLEX (Beckman). For Cas9-based MMEJ reporter assay using HEK293T *POLQ* WT or knockout cells, the MMEJ reporter, Cas9 and sgRNA targeting the reporter (co expressing BFP) together with a control vector, *POLQ* WT cDNA or *POLQ* mutant cDNA were transduced using Lipofectamine LTX with Plus reagent (Thermo Fisher Scientific). After 72 h incubation, GFP signals within BFP positive fraction were analyzed by CytoFLEX (Beckman).

### Comet assay

The alkaline comet assays were performed to detect both single and double-stranded DNA breaks. 1000 cells, suspended in PBS, were mixed with 50μl melted low melting agarose and pipetted onto the Cometslides (R&D Systems). Once the agarose solidified, the slides were immersed in a lysis solution for 18 hours to facilitate cell lysis. Following lysis, the slides were incubated in an alkaline unwinding solution for 1 hour to denature the DNA. Electrophoresis was performed at 21 V for 45 minutes in an alkaline electrophoresis solution. The slides were stained with SYBR green solution, scanned by fluorescence microscope, and analyzed using CometScore 2.0 software.

### Chromatin immunoprecipitation (ChIP) assay

U2OS biallelic endogenous V5-*POLQ* expressing cells carrying an Isce-I based MMEJ reporter were adenovirally transuduced with Isce-I in the presence or absence of 100ng/ml nocodazole. 12h after Isce-I trunscduction, chromatin immunoprecipitation was performed using SimpleChIP Enzymatic Chromatin IP Kit (Magnetic Beads) (Cell Signaling Technology). Protein-DNA cross-linking was performed with 1% formaldehyde and glycine solution. Chromatin was fragmented with Micrococcal Nuclease. The digested chromatin was incubated at 4°C with each antibody for 4 h. After the antibody reaction, ChIP-Grade Protein G Magnetic Beads were added to the samples, and the samples were incubated at 4°C for 1–2 h. For crosslinking reversal, each sample was incubated at 65°C with Proteinase-K overnight. After DNA purification, immunoprecipitated DNA was analyzed by qPCR using a primer set targeting the integrated MMEJ reporter locus. ChIP-qPCR was performed in technical triplicate. All the primer sequences for ChIP-qPCR used in this study were provided in Supplemental Table 1.

### Reverse transcription PCR (RT-PCR)

Total RNA was extracted using RNeasy Mini Kit (QIAGEN). Complementary DNA synthesis was performed using SuperScript IV First-Strand cDNA Synthesis Reaction (Thermo Fisher Scientific). The PCR products were separated in 2% agarose gel stained with SYBR Safe (Thermo Fisher Scientific) and imaged using ChemiDoc MP Imaging System (Bio-rad). All the primer sequences for RT-PCR used in this study were provided in Supplemental Table 1.

### ASO transfection

ASO was designed to target near the splice sites of the *POLQ* exon 25. The ASO was synthesized using 2ʹ-O-methoxyethyl-phosphorothioated bases by Integrated DNA Technologies (IDT). Prior to ASO transfection, cells were seeded in a 6-well plate and cultured overnight. ASO was transfected at a final concentration of 0.1 µM using Lipofectamine LTX with Plus reagent (Thermo Fisher Scientific). After 48 h incubation, cells were harvested and analyzed by RT-PCR and western blotting. All the sequences for ASO used in this study were provided in Supplemental Table 1.

### Splicing minigene reporter assay

A mini-gene reporter carrying mNeongreen split by synthetic intronic sequence derived from *POLQ* intron 24/25 and exon 25 was generated as previously described ^28^. The synthetic intron was made up of 250 bp of the start of the *POLQ* intron 24, 250 bp or 1000bp of the end of the *POLQ* intron 24, the *POLQ* exon 25, 250 bp or 1000bp of the start of the *POLQ* intron 25m and 250 bp of the end of the *POLQ* intron 25. The synthetic introns and exon so generated were placed between the sequences encoding the N-terminus and C-terminus of split mNeongreen. The mScarlet-P2A-split mNeongreen (N-terminus)-synthetic introns/exon-split mNeongreen (C terminus) construct was integrated into a pLV-EF1a-IRES-Neo vector (Addgene #85139) to generate the mini-gene reporter. K562 cells or Ewing sarcoma cell lines were lentivirally transduced with the mini-gene reporter. 48hr after lentiviral transduction, the cells were enriched by using G418 (0.8mg/ul) for seven days. After antibiotic selection, mScarlet and mNeongreen signals in the reporter cells were analyzed by CytoFLEX (Beckman). For the flow cytometry analyses, mScarlet positive fractions within living cells were gated. Thereafter, the frequency of the mNeongreen positive cells within mScarlet expressing cells was analyzed by using Flowjo ver.10.9.0.

### Mass spectrometry analysis

HEK293T EWSR1-FKBP12 V36F knock-in cells were seeded into 15cm dishes and cultured overnight. The cells were transfected with pMYS-IB empty vector (EV) or pMYs-IB-EWSR1 wildtype or pMYs-IB-EWSR1 37YS plasmid. Cells were transfected with 30 µg of vector /dish using Lipofectamine LTX with Plus reagent. 16h after transfection, media was changed with 1uM dTAG^v1^. The cells were harvested 40 h after transfection and then were lysed with 40 mM Tris-HCl pH 8 buffer supplemented with 0.5% NP-40, 200 mM NaCl, 2 mM EDTA, Protease/Phosphatase inhibitor Cocktail (Cell Signaling Technology), and MG-132. The lysate was centrifuged, and the pellet was resuspended with 20 mM Tris-HCl pH 7.5 supplemented with 100 mM KCl, 2 mM MgCl2, 1 mM CaCl2, and Protease/Phosphatase inhibitor Cocktail and treated with Micrococcal Nuclease (Thermo Fisher Scientific) for approximately 30 minutes, followed by addition of 5 mM EGTA (final concentration) to stop digestion. After three pulses of sonication (30 s each), samples were centrifuged to collect the supernatants as chromatin fractions. Immunoprecipitation was performed using Anti-FLAG M2 antibody (Sigma-Aldrich) conjugated with Dynabeads Protein G for Immunoprecipitation (Thermo Fisher Scientific). Beads were washed with TGN-150 wash buffer containing 20 mM Tris-HCl pH 7.6, 150 mM NaCl, 3 mM MgCl2, 10% glycerol, and 0.01% NP-40. Beads were then boiled at 70nC for 20 minutes in 1:1 diluted NuPAGE LDS Sample buffer (4X) (Thermo Fisher Scientific) to elute the proteins. The eluted proteins were applied to SDS-PAGE, and Coomassie stained gel band samples were used for the analyses. Two independent biological replicate samples were analyzed by mass spectrometry.

The gel bands were cut into approximately 1 mm3 pieces. Gel pieces were then subjected to a modified in-gel trypsin digestion procedure. Gel pieces were washed and dehydrated with acetonitrile for 10 minutes and completely dried in a SpeedVac (Thermo Fisher Scientific). The gel pieces were rehydrated with 50 mM ammonium bicarbonate containing 12.5 µg/mL modified sequencing-grade trypsin (Promega) at 4°C. After rehydration, the excess trypsin solution was removed and replaced with 50 mM ammonium bicarbonate solution. Samples were then incubated at 37°C overnight. Peptides were later extracted with a solution containing 50% acetonitrile and 1% formic acid and dried in a SpeedVac. The samples were reconstituted in 5–10 µL of HPLC solvent A (2.5% acetonitrile, 0.1% formic acid). A nano-scale reverse-phase HPLC capillary column was created by packing 2.6 µm C18 spherical silica beads into a fused silica capillary (100 µm inner diameter x ∼30 cm length) with a flame-drawn tip. After equilibrating the column each sample was loaded via an EASY-nLC (Thermo Fisher Scientific). A gradient was formed, and peptides were eluted with increasing concentrations of solvent B (90% acetonitrile, 0.1% formic acid). As peptides eluted, they were subjected to electrospray ionization and then entered into a Orbitrap Exploris480 mass spectrometer (Thermo Fisher Scientific). Peptides were detected, isolated, and fragmented to produce a tandem mass spectrum of specific fragment ions for each peptide. Peptide sequences were determined by matching protein databases with the acquired fragmentation pattern by the software program, Sequest (Thermo Fisher Scientific). All databases include a reversed version of all the sequences and the data was filtered to between a one and two percent peptide false discovery rate. Perseus (version 2.0.11.0) ^69^ was used to preprocess the protein data. The protein intensities were normalized by the median and log2-transformed. Proteins with only one intensity value or with a sum of all total peptide counts of less than 5 were filtered out. For each sample, missing log2-intensity values were imputed using the normal distribution with width of 0.3 and down shift of 1.8. Histograms of log2-instensity values were inspected to confirm that the imputed values were at the lower end. Finally, limma analysis (version 3.54.2) ^70^ was performed on the log2-instensity values to determine enriched and depleted proteins. Volcano plots were generated using ggplot2 and ggrepel R packages.

### RNA immuno-precipitation (RIP) assay

Protocol for RIP assay was adapted from Nicholson-Shaw et al., 2022^29^. HEK293T EWSR1-FKBP12 V36F knock-in cells were cultured in 15 cm dishes and 2-3 plates were used per cell line for each assay. 16hr after transfection, media was changed with 1uM dTAG^v1^. Cells were harvested 40hr after transfection, washed with ice-cold PBS 2X times, cross-linked with 0.2% PFA for 15 min. and then quenched by adding glycine to a final concentration of 125mM. Cells were lysed in iCLIP buffer (50mM Tris-HCl pH 7.4, 100mM NaCl, 1% NP-40, 0.1% SDS, 0.5% sodium deoxycholate) supplemented with protease, phosphatase and RNase inhibitor cocktail. The samples were rotated at 4°C for 10 min. and then sonicated on ‘light’ setting twice. The lysate was incubated with Protein A beads on a rotor for 10 min. at 4°C to pre-clear the extract. The lysate was centrifuged at max speed for 10 min. and the supernatant was recovered and incubated for 2 hrs on a rotor at 4°C with Protein A beads coupled to EWSR1 Ab (Life Technologies). A part of the lysate (0.4-0.6%) was kept aside as input. After 2 hrs the beads were washed twice with iCLIP buffer. The input and the IP samples were all subjected to TURBO DNase treatment (Thermo Fisher Scientific) for 30 min. at 37°C to get rid of the genomic DNA and the cross-links were reversed by digesting with Proteinase K (NEB) for 30 min. at 37°C and further digestion and denaturation of proteins was achieved by adding urea to a final concentration of 2.5M and incubating the samples at 37°C for additional 20 min. After digestion of the DNA and protein from the samples, the RNA was purified by adding Trizol and processing the samples through Directzol RNA mini-prep kit (Zymo Research). The eluted RNA from the input and IP samples were converted to cDNA using the High-capacity RNA-to-cDNA kit (Thermo Fisher Scientific). The reverse transcription was performed with and without adding the reverse transcriptase enzyme. qPCR was set up with primers targeting *POLQ* pre-mRNA, *POLQ* mature mRNA, and GAPDH in technical replicates for each assay. Ct values were normalized to that from the control samples. . All the sequences for RIP-qPCR used in this study were provided in Supplemental Table 1.

### Next generation sequencing

Library preparation and sequencing reactions were conducted at GENEWIZ, Lin/Azenta US, Inc. Deep RNA-seq (300 million reads/sample, N = 3 for each cell line) was performed for RPE *p53-/-* (parental control) and *p53-/-EWSR1* knock-out cells. RNA samples were prepared using RNeasy Mini Kit and RNase-Free DNase Set (QIAGEN). RNA samples were quantified using Qubit 2.0 Fluorometer, and the RNA integrity was checked with 4200 TapeStation. NEBNext Ultra II RNA

Library Prep Kit for Illumina was used following the manufacturer’s recommendations (New England BioLabs). Briefly, mRNAs were initially enriched with Oligo d(T) beads. Enriched mRNAs were fragmented for 15 minutes at 94°C. First strand and second strand cDNA were subsequently synthesized. cDNA fragments were end repaired and adenylated at 3ʹends, and universal adapters were ligated to cDNA fragments, followed by index addition and library enrichment by PCR with limited cycles. The sequencing libraries were validated using Agilent TapeStation and quantified using Qubit 2.0 Fluorometer as well as by quantitative PCR (KAPA Biosystems). The sequencing libraries were clustered on flowcell lanes. After clustering, the flowcell was loaded on the Illumina NovaSeq instrument according to manufacturer’s instructions. The samples were sequenced using a 2×150 Paired End configuration. Image analysis and base calling were conducted by the Control Software (NCS). Raw sequence data (.bcl files) generated from the Illumina instrument was converted into fastq files and de-multiplexed using Illumina’s bcl2fastq (version 2.17) software. One mismatch was allowed for index sequence identification. Several bioinformatics analyses were performed on the RNA-seq data to identify differentially expressed genes, differentially used exons, and differentially spliced events between RPE *p53*-/-control cells and *p53-/-EWSR1* knock-out cells.

For differential gene expression analysis, Kallisto (version 0.48.0) ^71^ was used to map the raw RNA-seq reads against the GENCODE human reference gene annotation (Release 44) ^72^ and generate raw transcript abundance estimates, and the *tximport* Bioconductor R package ^73^ was used to summarize the transcript abundance estimates to produce gene-level estimated counts. Differentially expressed genes between control and *EWSR1* knock-out cell samples (N = 3) were determined using both edgeR (version 3.40.2) ^74^ and DESeq2 (version 1.38.3) ^75^ Bioconductor packages. The high concordance between the estimated log_2_ foldchange of the two algorithms was verified.

For both differential exon usage and differential splicing analyses, STAR (version 2.7.10b) ^76^ was used to map RNA-seq reads and generate BAM files. Specifically, the 2-pass STAR was used to map the raw RNA-seq reads against the GRCh38/hg38 human reference genome, with the GENCODE human reference gene annotation (Release 44) ^72^. The first round of the 2-pass STAR was run to identify splice junctions, and more confidence junctions then were filtered by requiring them to have at least 5 total mapped reads. The second round of the 2-pass STAR was run using the filtered junctions to generate BAM files.

For differential exon usage analysis, DEXSeq (version 1.46.0) ^77^ was used. In brief, a list of unique exonic regions was initially generated using the Python script *dexseq_prepare_annotation.py.* This script “collapsed” exonic regions from different transcripts to be used as exon counting bins. Next, the *dexseq_count.py* Python script was used to count reads for each sample. Finally, differentially used exons between control and *EWSR1* knock-out cells (N = 3) were determined using DEXSeq. For the differential splicing analysis, MISO (version 0.5.4) ^26^ and other tools were used as discussed below. The exon-centric splicing event annotations were generated using the *rnaseqlib* Python package (http://github.com/yarden/rnaseqlib). These annotations were derived considering all transcripts from Ensembl, UCSC RefSeq and NCBI RefSeq gene annotations, which were obtained from https://hgdownload.soe.ucsc.edu/goldenPath/hg38/database, in files *knownGene.txt*, *refGene.txt*, and *ncbiRefSeq.txt*. Each annotation event belongs to one of these 5 types: skipped exons (SE), alternative 3ʹ splice sites (A3SS), alternative 5ʹ splice sites (A5SS), mutually exclusive exons (MXE), and retained introns (RI). These annotated events were linked to representative overlapping exons or introns of the GENCODE human reference gene annotation (Release 44) using the intersect function of the *bedtools* toolset (version 2.31.0). The representative exons and introns were prioritized with a high overlapping percentage (95%), followed by good transcript quality information including whether they were annotated as canonical and basic, and with better transcript support level. MISO was performed to quantify the isoforms and compute percent-spliced-in values (PSIs) of each alternative splicing event from the sorted BAM of each sample. Lower quality events were filtered out as follows. For each replicate sample pair, the *compare_miso* program was used to compare the results of the control and the *EWSR1* knock-out samples. The *filter_events* program was then used to filter the comparison results, by requiring each sample to have at least 35 total isoform-identifying reads. Finally, *limma* analysis (version 3.54.2) was performed on the PSI values of the filtered events to determine differential splicing events between control and *EWSR1* knock-out cells (N = 3). Significant differential alternative splicing events were called if they had *FDR* smaller than 0.05 and absolute delta PSI of at least 0.05. Events that had significant FDR (<0.05) but small absolute delta PSI (<0.05) were likely to be artifacts and thus were excluded from *limma* results. Volcano plots were generated using ggplot2.

To have more confidence in the MISO results, we also independently analyzed differential splicing using rMATS (version v4.2.0) ^27^. Since SE events were the most frequent event type, we focused on analyzing the concordance of SE events between MISO and rMATS. First, rMATS was run using all sorted BAMs as inputs and the GENCODE human reference gene annotation (Release 44). Lower quality events were filtered out by requiring each sample to have at least 35 total isoform-identifying reads. Then, the shared events that were highly similar between MISO and rMATS were identified if their target regions overlapped by at least 95%.

Since one MISO event could be linked to multiple slightly different rMATS events, for each MISO event, a representative rMATS event was chosen that had the largest absolute difference in inclusion level. Finally, we assessed the concordance between MISO and rMATS by comparing their inclusion differences (delta PSI values and differences in inclusion levels, respectively) from these shared events. We observed that a great majority of significant MISO events had a shared rMATS event. Furthermore, we observed that the inclusion differences of MISO and rMATS were positively correlated with R^2^ of 0.32. The scatterplot comparing the inclusion differences of the two algorithms was generated using ggplot2. Sashimi plots were generated using the *rmats2sashimiplot* program (https://github.com/Xinglab/rmats2sashimiplot/).

### AlphaFold analysis

AlphaFold3 and the AlphaFold Server ^78^ were used to model full-length (POLQ aa 1824-2579) and truncated (POLQ aa 1824-2390) POLQ polymerases domains in complex with two complementary 9 bp DNA molecules (5’-CCAATGACA-3’ and 5’-TGTCATTGG-3’). Models were visualized in Pymol V3.0.4 (Schrödinger) using the top-ranking predictions per seed. The full-length model was compared with a published crystal structure of the POLQ polymerase domain (PDB 6XBU) ^79^ to ensure its accuracy.

### Mouse xenograft study

All animal experiments were conducted in accordance with Institutional Animal Care and Use Committee-approved protocol (#08-036) at Dana-Farber Cancer Institute. TC32 cells were lentivirally transduced with luciferase by using a pLenti CMV Puro LUC (w168-1) vector. Two million TC32 cells expressing luciferase were subcutaneously injected into 7-week-old male and female NOD.Cg-*Prkdc*^scid^ Il2rg^tm1Wjl^/SzJ mice. After confirming tumor engraftment, mice were treated with Peposertib (100 mg/kg) or vehicle control once a day via oral administration together with Etoposide (8mg/kg) or vehicle control via intraperitoneal injection (IP injection) twice a week for 2 weeks. Tumors were measured every 2 to 3 days using an electronic caliper, and tumor volumes were calculated by using the formula L×W×W/2. For immunohistochemistry analysis, tumors were excised and fixed with formaldehyde. Mice with a tumor of more than 20 mm in length or width were euthanized.

### Data acquiring and processing of transcriptomic sequencing from patient tumor samples

Patient tumor sample bulk transcriptomic sequencing data were obtained from St. Jude’s Cloud ^80^. For expression analysis, we used the mapped expression counts data; for the exon splicing data, we used the BAM files provided. For the expression counts data, all of the samples were uniformly processed and mapped to the same reference genome as part of the St. Jude’s Cloud resource, and as such, no correction between samples or cancer types was performed. To evaluate the *POLQ* expression, we first computed the TPM (transcript-per-million) counts for all samples. In this computation, we omitted genes annotated as pseudogenes in GENCODE v.31, and used the median gene length as the gene length for each gene symbol.

### Computation of exon skipping rates in primary patient samples

To compute the proportions of reads with exons skipped in the sequencing reads data obtained from St. Jude’s Cloud ^81^, rMATS-turbo, a computational tool to quantify alternative splicing patterns from RNA sequencing data, was applied. To compute the proportion of reads that each exon was skipped in, the rMATS-turbo results inclusive of supporting reads from counted from both junction and exon reads were used. Of note, for each individual exon, any sample that had no supporting reads for both the inclusive and skipped cases was omitted. All comparisons between distributions of skipped read proportions were performed using Mann-Whitney U tests.

### Mutational Analysis of Tumors

Mutational analysis was performed on the whole genome sequencing (WGS) data of Ewing sarcoma and neuroblastoma tumors obtained from St. Jude Cloud (https://www.stjude.cloud/). The samples include 23 Ewing sarcoma (ES) and 48 neuroblastoma (NB) tumor and matched-normal pairs. Somatic variants were determined as follows. First, for each cohort, short variants were called from the HG38 Genomic Variant Call Format (gVCF) data using the GATK joint genotyping workflow (version 4.2.4). Second, bcftools (version 1.14) was used to filter out low-quality variants with these hard filters: QD < 7.5, MQ < 59.9, -g3, and -G10. Only biallelic SNPs and INDELs were selected. Third, cohort-level singleton variants were selected if there was only one sample with read depth (DP) between 17 and 300, and variant allele fraction (VAF) of at least 0.1. And finally, a variant was considered somatic if both the tumor and matched-normal had SOR <= 3, MQ > 59.9, and DP > 17, and either, tumor VAF >= 0.175 and matched normal VAF = 0, or tumor VAF <= 0.825 and matched normal VAF = 1.

Profiles of SNV mutations and INDELs were generated using SigProfilerMatrixGenerator (version 1.2.13; https://github.com/AlexandrovLab/SigProfilerMatrixGenerator). Mutational signature analysis was performed using SigProfilerAssignment (version 0.0.21; https://github.com/AlexandrovLab/SigProfilerAssignment). This involved fitting the mutational profiles to the COSMIC (https://cancer.sanger.ac.uk/signatures/) SNV signatures (SBS version 2) and INDEL signatures (ID version 3.3).

The MMEJ signature was characterized by the presence of microhomologies at the boundaries of deletions and the presence of nearby template sequences in insertions, or templated insertions ^82,83^. The presence of microhomologies in deletions was also independently determined using the method described by Taheri-Ghahfarokhi *et al* ^82,83^. This method essentially finds the longest possible complementary subsequence between one end of the deleted sequence and the flanking sequence at the other end. In brief, for each deletion, 50-bp flanking sequences around the deleted sequence were first retrieved from the reference human genome (HG38) using bedtools (version 2.27.1; https://bedtools.readthedocs.io/). The deletion was then repeatedly shifted leftward by one base whenever its rightmost base matched the reference base immediately to its left. Finally, the microhomology sequence was the longest matching subsequence between the deleted sequence and the right flanking sequence starting from their leftmost bases. Deletions with length of at least 3bp with GATK ReadPosRankSum of greater than -5 and with microhomology of at least 2 bp were designated as MMEJ events ^82^. For comparison between ES and NB, tumors with at least 15 deletions and the proportions of MMEJ deletions among all indels were used. Local templated insertions were characterized by the presence of subsequences in the inserted sequence that were identical to the flanking sequences (templated inserts) ^15,84,85^. The templated inserts were detected by aligning the inserted sequence with the 50bp-flanking sequences around the insertion site using the Smith–Waterman local alignment algorithm ^86^. The analysis was restricted to insertions with GATK ReadPosRankSum of greater than -5 and with length of at least 5bp. The insertions with templates adjacent to the insertion position were excluded. The remaining insertions with templates covering at least 60% of the inserted sequence were designated as MMEJ events. For comparison between ES and NB, tumors with at least 2 insertions, and the proportions of MMEJ templated insertions among all insertions were used.

### Immunohistochemistry (IHC)

Primary patient samples for IHC analysis were obtained from the Mayo Clinic (IRB approval #24-007702). IHC was performed using a Parhelia autostainer. Slides were deparaffinized using Slide Brite xylene substitute (23401-01) followed by rehydration in ethyl alcohol. Endogenous blocking was performed using methanol hydrogen peroxide solution. Slides were heated for 30 minutes in citrate solution for epitope retrieval. Slides were incubated for 30minutes in anti-DNAPKcs phosphor S2056 antibody at 1:150 followed by washing in wash buffer (Dako S3006) and inbubation for 30minutes in Mach 3 rabbit HRP (Buocare Medical RP531L). Primary staining was performed by incubation in DAB (Dako K3468) for 10minutes. Counterstaing was performed using hematoxylin (Sigma-Aldrich MHS16) and bluing buffer (Dako C5702). Slides were then dehydrated in ethyl alcohol before coverslipping using Permount mounting mediun (Fisher Scientific SP15-100). Slides were digitally imaged using a Motic Easyscan Pro 6.

Histological image analysis was performed using QuPath (version 0.5.1). Whole-slide images (WSIs) were imported into QuPath, and automated color deconvolution was applied to separate the hematoxylin and DAB (3,3′-diaminobenzidine) staining channels. The mean optica l density (OD) of the DAB-positive areas was measured to assess staining intensity. For patients with multiple tissue sections, and the mean OD across all sections for the same patient was used to represent the patient’s staining intensity.

### Statistical analysis

Data were analyzed and visualized using GraphPad Prism (Version 10.2.2, GraphPad Software, LLC). For comparisons between two groups, either a two-tailed Student’s t-test or a Mann-Whitney U test was used. For comparisons involving more than two groups, one-way analysis of variance (ANOVA) was performed. For *in vivo* survival analysis study, a log-rank test was performed. A p-value of less than 0.05 (p < 0.05) was considered statistically significant.

## ACKNOWLEDGMENTS

We thank members of the D’Andrea laboratory for their helpful suggestions and comments. We thank Harvard Medical School and the Dana-Farber/Harvard Cancer Center in Boston, MA, for the use of the Animal, Flowcytometry, and Taplin Mass Spectrometry core facilities. This work was supported by U.S. National Institutes of Health grants R01HL052725 and R01CA296618 (A.D.D), the Breast Cancer Research Foundation (A.D.D.), the Ludwig Center at Harvard (A.D.D.), the David Liposarcoma Research Initiative at Harvard (A.D.D.), and the Smith Family Foundation (A.D.D.), and the Alex’s Lemonade Stand Foundation (S.A.). This work was also supported by U.S. National Institutes of Health grants P50 CA168504 (A.D.D, G.I.S.), P50 CA240243 (A.D.D., G.I.S.), the SENSHIN Medical Research Foundation (S.A.), the Japan Society for the Promotion of Science Overseas Research Fellowships (S.A.), NCI K08 Career Development Award (R.G.), the DoD PRCRP Fellow Career Development Award (R.G.), the ASCO Career Development Award (R.G.), the Rally Foundation Career Development Award (R.G.) and the Boston Children’s Hospital TRP Mentored Career Development Award (R.G.).

## AUTHOR CONTRIBUTIONS

S.A and A.D.D. conceived the study. S.A., G.Z., J.P.A., Y.H., D.R.I., S.M. and L.J. designed and performed biological experiments and analyzed the data. S.A. and Y.H designed and performed animal experiments. Y.T., H.N. and R.G. performed computational analysis. N.W.A. performed AlphaFold analysis. M.D.B., J.J.T and S.I.R. performed IHC analysis using patient samples. K.P., E.M.V.A. and G.I.S. provided scientific inputs. S.A. and A.D.D. wrote the manuscript.

---

**Supplementary Figure 1:**
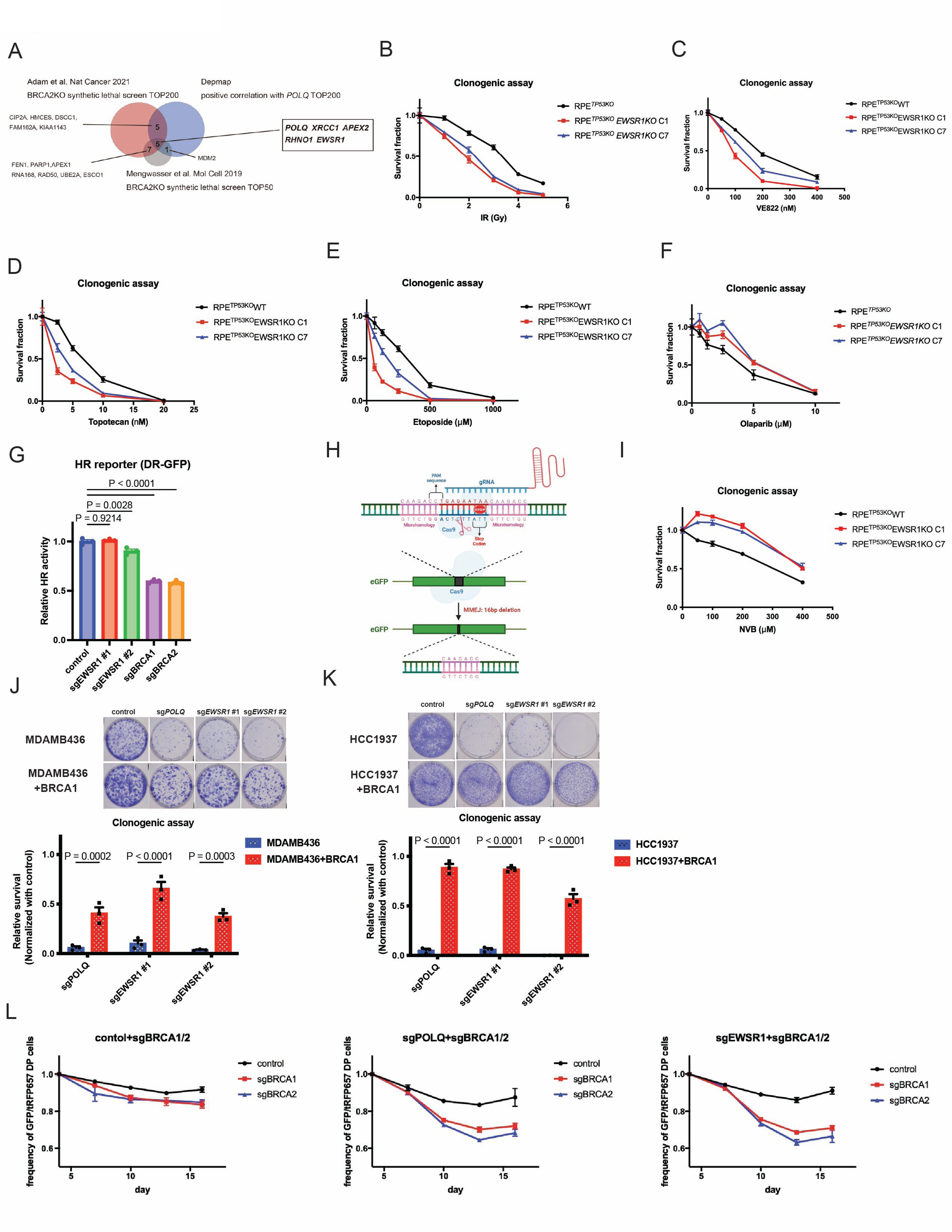
*EWSR1* loss is synthetic lethal with BRCA1/2 deficiency. (A) A Venn diagram showing overlap between genes identified in the previously reported CRISPR screens to identify synthetic *BRCA2* lethality and genes whose dependence is positively correlated with *POLQ* in the Depmap analysis. (B-F, I) Colony formation assay data of RPE *p53*-/-cells and RPE *p53*-/-*EWSR1* knock-out clones treated with X-ray (B), VE822 (C), Topotecan (D), Etoposide (E), Olaparib (F) and Novobiocin (I 4) (n=4). (G) U2OS DR-GFP reporter cells (I-SceI based) were transduced lentivirally transduced with control sgRNA or sgRNA targeting EWSR1, BRCA1 or BRCA2 (co-expressing puromycin resistance gene). Cells were treated with puromycin for 4days, and then subjected to reporter analysis (n=3). (H) Schematic representation of the Cas9-based MMEJ reporter assay (ver.2). (J, K) Colony formation assay data of MDA-MB-436 (J, n=3) and HCC1937 (K, n=3) cells and the cells complemented with BRCA1 cDNA transduced with control, sg*POLQ* or sg*EWSR1* (left). Shown are representative images of the colonies (right). (L) RPE *p53*-/-cells were lentivirally transduced Cas9 (expressing balasticidin resistance gene). After balasticicin selection, cells were lentivirally transduced with a sgRNA targeting control, *POLQ* or *EWSR1* (co-expressing GFP) together with with a sgRNA targeting control, *BRCA1* or *BRCA2* (co-expressing tRFP657). 72hr after the sgRNA transduction, the frequency of the GFP/tRFP657 cells was traced by FACS (n=3).

**Supplementary Figure 2:**
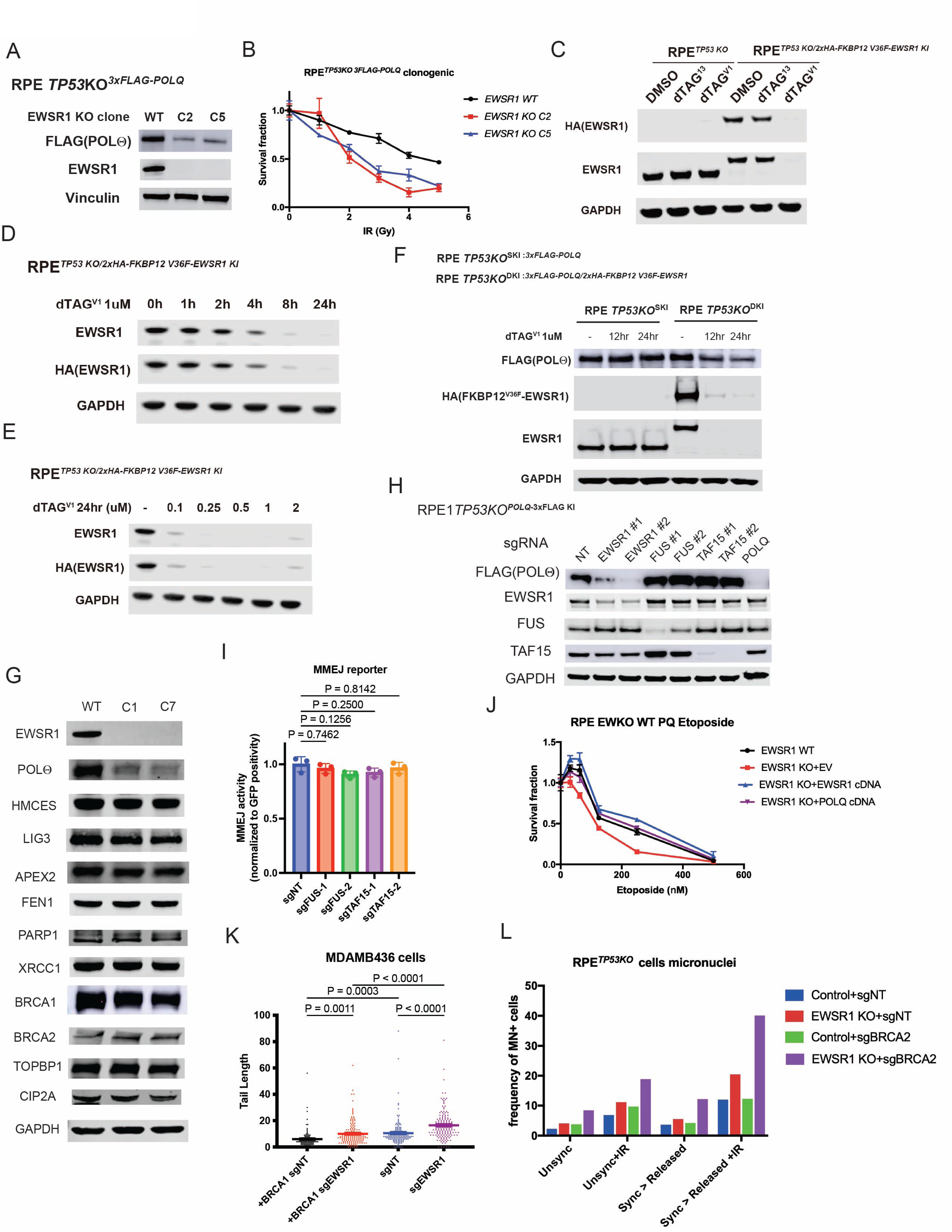
Subacute depletion of EWSR1 results in POL0 protein loss. (A) RPE *TP53-/-* 3xFLAG-POLQ knock-in cells were transduced with Cas9 and a sgRNA targeting *EWSR1* by electroporation. Single clones are isolated 7days after electroporation. Shown are immunoblotting analyses for FLAF-POL0, EWSR1 and GAPDH in control cells and *EWSR1* knockout clones. (B) Colony formation assay data of RPE *p53*-/-3xFLAG-*POLQ* knock-in cells and RPE *p53*-/-*EWSR1* knock-out 3xFLAG-*POLQ* knock-in clones treated with X-ray (n=3). (C) RPE *p53*-/-2xHA-FKBP12 V36F *EWSR1* cells were treated with DMSO, 1μM dTAG^13^ or 1μM dTAG^v1^ for 24hr. The cell lysates were extracted and then analyzed by immunoblotting. dTAG^v1^ but not dTAG^13^ efficiently degraded endogenous EWSR1 protein. (D) RPE *p53*-/-3xFLAG-*POLQ*/2xHA-FKBP12 V36F *EWSR1* double knock-in cells are treated with 1μM dTAG^V1^ for the indicated time. The cell lysates were extracted and then analyzed by immunoblotting. (E) RPE *p53*-/-3xFLAG-*POLQ*/2xHA-FKBP12 V36F *EWSR1* double knock-in cells are treated with dTAG^V1^ at the indicated concentration for 24hr. The cell lysates were extracted and then analyzed by immunoblotting. (F) RPE *p53*-/-3xFLAG-*POLQ* single knock-in cells and RPE *p53*-/-3xFLAG-*POLQ*/2xHA-FKBP12 V36F *EWSR1* double knock-in cells are treated with dTAG^V1^ for indicated time. The cell lysates were extracted and then analyzed by immunoblotting. (G) Immunoblotting analyses showed that BRCA1, BRCA2 and other MMEJ proteins, except for POL0, were not affected by *EWSR1* knockout in RPE *p53*-/-cells. (H) RPE *p53*-/-3xFLAG-*POLQ* knock-in cells were lentivirally transduced with control sgRNA or sgRNA targeting *EWSR1, FUS* or *TAF15* (co-expressing puromycin resistance gene). Cells were treated with puromycin for 4days, and then the cell lysates were extracted and then analyzed by immunoblotting. (I) K562 MMEJ ver.2 (Cas9-based) cells were lentivirally transduced with control sgRNA or sgRNA targeting *EWSR1, FUS* or *TAF15* (co-expressing puromycin resistance gene). Cells were treated with puromycin for 4days, and then cells were subjected to MMEJ reporter analysis (n=3). (J) RPE *p53*-/-cells and RPE *p53*-/-*EWSR1* knockout cells were transduced with control, wildtype *EWSR1* or *POLQ* cDNA. Colony formation assays were performed in the presence of Etoposide (n=3). (K) MDA-MB-436 cells and the cells complemented with BRCA1 were transduced with Cas9 and a sgRNA targeting control or *EWSR1*. Cells were subjected to comet assay to measure unrepaired DNA damage. (L) RPE *p53*-/-cells and RPE *p53*-/-*EWSR1* knockout cells were transduced with Cas9 and a sgRNA targeting control or *BRCA2*. Cells were treated with or without synchronization by 8μM RO3306. 1 hour after removal of RO3306, cells were treated with 2 Gy irradiation and then incubated for 24 hours. The frequency of micronuclei positive cells is shown.

**Supplementary Figure 3:**
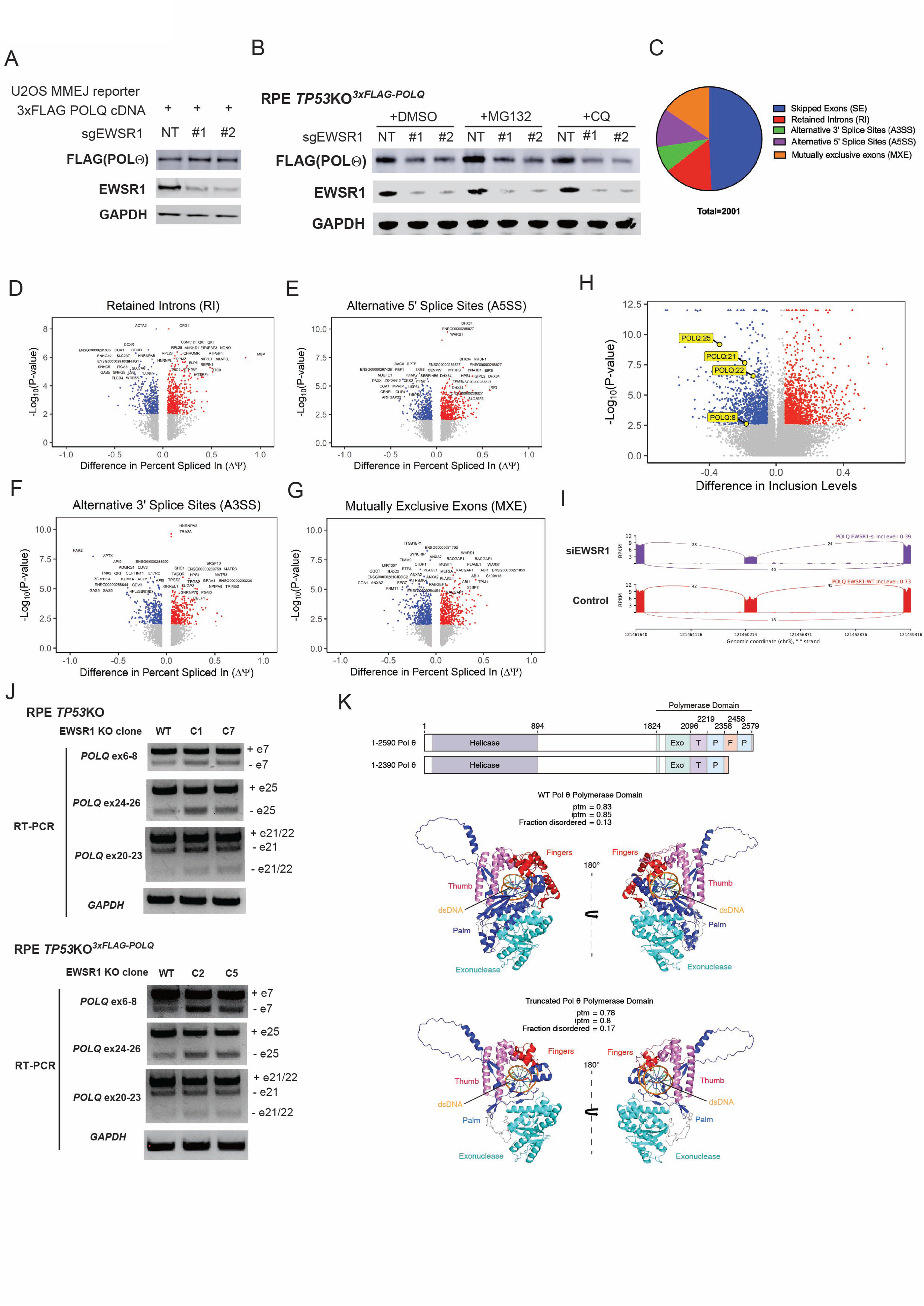
EWSR1 prevents multiple exon skipping events of the *POLQ* transcript. (A) U2OS MMEJ reporter cells (I-SceI based) were transduced with *POLQ* cDNA (co-expressing blasticidin resistance gene). After blasticidin selection, cells were subsequently transduced with Cas9 and a sgRNA targeting control or *EWSR1* (co-expressing puromycin resistance gene). After puromycin selection, cell lysates were extracted and then analyzed by immunoblotting. See also Fig.2C. (B) RPE *p53*-/-3xFLAG-*POLQ* knock-in cells were transduced with Cas9 and a control sgRNA or sgRNA targeting *EWSR1* by electroporation. 72hr after electroporation, cells were treated with MG132 or Chloroquine for 4hr. After the treatment, cell lysates were extracted and then analyzed by immunoblotting. (C) A pie chart summarizing alternative splicing event types in RPE *p53*-/-*EWSR1* knock-out cells obtained by MISO analysis. (D-G) Volcano plots showing retained introns (RI) (D), alternatively spliced 5’ splice cites (A5SS) (E), alternatively spliced 3’ splice cites (A3SS) (F), mutually exclusive exons (MXE) (G) in RPE *p53*-/-*EWSR1* knock-out cells compared with RPE *p53*-/-cells. (H) Volcano plots showing alternatively spliced skipped exon (SE) in HEK293T si*EWSR1* transduced cells compared with control cells defined by RNA-seq analysis (GSE154944). (I) Sashimi plots of the alternative splicing events in control and *EWSR1* knock-out RPE *p53*-/-cells: the region between exon 23 and exon 26 of *POLQ* transcript. (J) RT-PCR showing the exon 7, 21, 22 and 25 skipping of *POLQ* transcript in *EWSR1* knock-out cells derived from RPE *p53*-/-(top) and from RPE *p53*-/-3xFLAG-*POLQ* knock-in cells (bottom). (K) The schematics illustrates the location of protein domains with the primary sequence POLΘ. Numbers refer to amino acid residues. Exo = exonuclease, T = thumb, P = palm, F = fingers. The ribbon illustrations are AlphaFold 3-generated predictions of the WT and truncated form of the POL0 polymerase domain. The two views represent 180° rotation around the y-axis. Colors illustrate the sub-domain architecture.

**Supplementary Figure 4:**
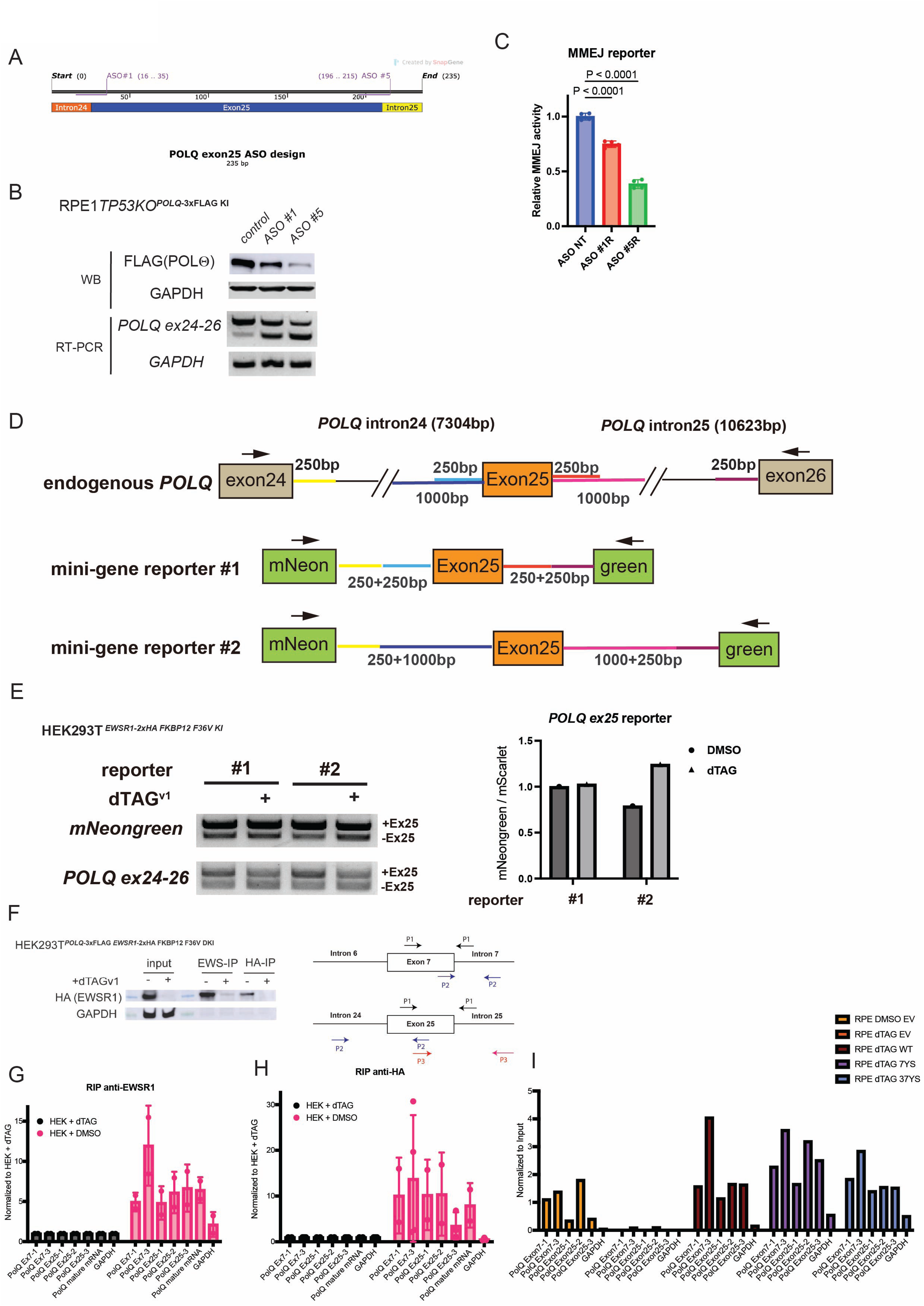
Exon 25 skipping of the *POLQ* transcript decreased POL0 protein, resulting in profound MMEJ deficiencies. (A) Schematics of the *POLQ* exon 25, intron 24/25 and the design of ASO and sgRNAs targeting the splice sites. (B) Immunoblotting (upper) and RT-PCR (lower) analysis showing that transduction of two ASOs efficiently induced exon 25 skipping and POL0 protein loss in RPE *p53*-/-3xFLAG-*POLQ* knock-in cells. (C) U2OS MMEJ ver.2 (Cas9-based) were transduced control or two ASOs targeting *POLQ* exon 25 splice sites. 48h after ASO transduction, cells were subjected to MMEJ reporter analysis (n=4). (D) Schematic representation of the endogenous *POLQ* exon 25 and intron 24/25 and the minigene constructs. (E) HEK293T 2xHA-FKBP12 V36F *EWSR1* dknock-in cells were transfected with the indicated minigene construct and then treated with DMSO or dTAG^V1^. 48h after transfection, cells were subjected to assess the endogenous-and minigene-derived *POLQ* exon 25 skipping (left) and FACS analysis to measure the reporter activity (right). (F-H) Binding of EWSR1 to *POLQ* pre-mRNA evaluated using the RNA immuno-precipitation (RIP) assay, performed using an anti-EWS antibody and an anti-HA antibody (N=2). The assay was performed in HEK293T 3xFLAG-POLQ/2xHA-FKBP12 V36F *EWSR1* double knock-in cells in the presence of DMSO or dTAG^v1^. qPCR was performed with primers targeting *POLQ* pre-mRNA around exon 7 and 25, *POLQ* mature mRNA and *GAPDH* (bottom). Shown were immunoblotting for cell lysates with or without immunoprecipitation (F), RIP-qPCR assay for the samples with EWS-IP (G) and HA-IP (H). (I) Binding of multiple EWSR1 mutants to *POLQ* pre-mRNA evaluated using the RNA immuno-precipitation (RIP) assay, performed using anti-EWS antibody. The assay was performed in HEK293T 3xFLAG-*POLQ*/2xHA-FKBP12 V36F *EWSR1* double knock-in cells transduced with a control vector, EWSR1 WT, EWSR1 7YS mutant or EWSR1 37YS mutant in the presence of DMSO or dTAG^v1^. qPCR was performed with primers targeting *POLQ* pre-mRNA around exon 7 and 25 and *GAPDH* (bottom).

**Supplementary Figure 5.**
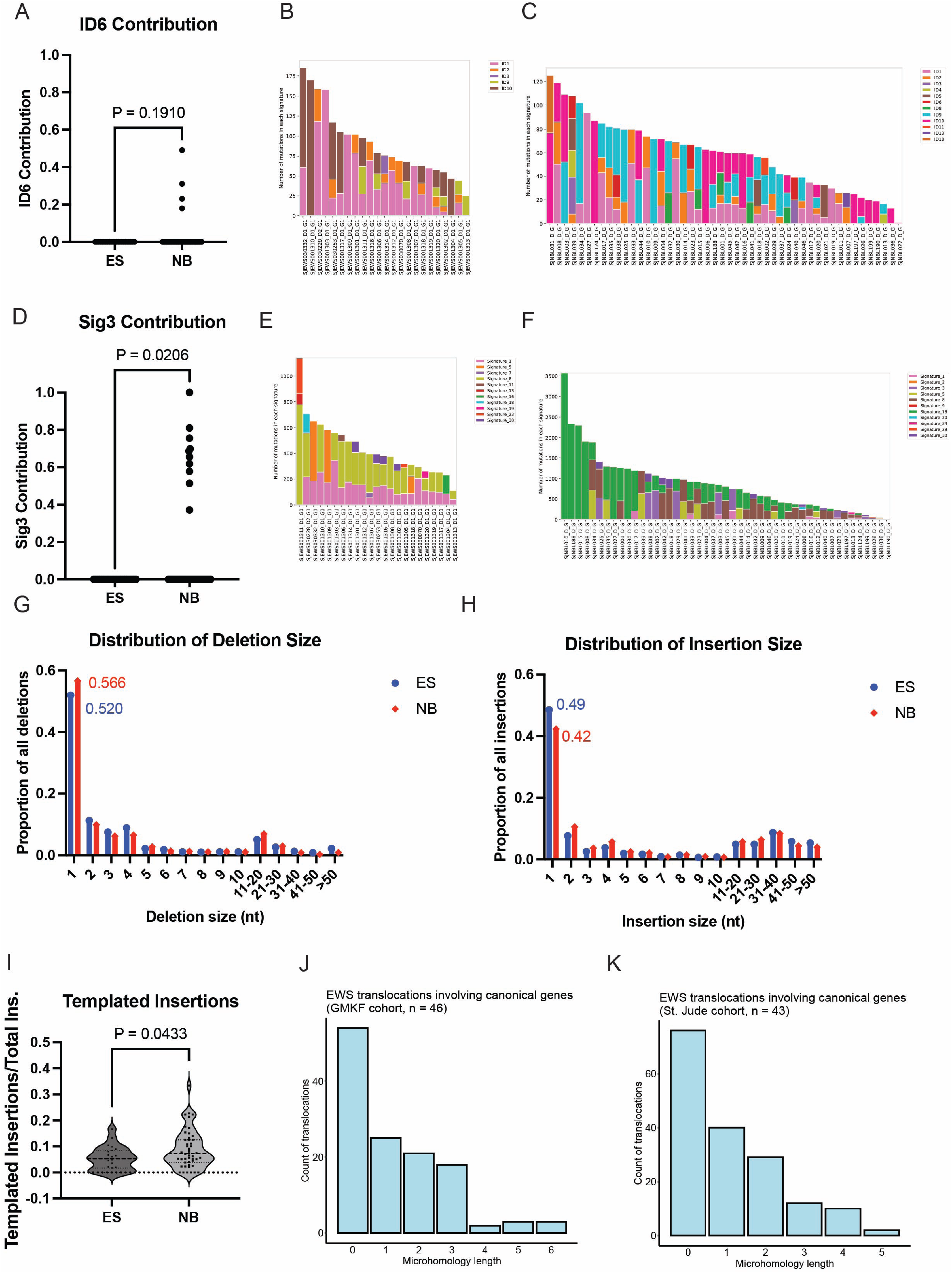
Samples from Ewing sarcoma patients show no HRD genomic signatures. (A-C) Contribution of COSMIC indel (ID)-signature (A) and stacked bar plots of the numbers of indels attributed to COSMIC ID signatures for Ewing sarcoma tumors (ES) (in B), and neuroblastoma tumors (NB) (in C). Short somatic mutations were determined from whole genome sequencing data of tumors and matched normals using GATK4, and SigProfilerAssignment was used to estimate the COSMIC ID signatures from the mutations. Each bar contains relative numbers of indels attributed to specific ID signatures, which are shown in different colors. There are no ES tumors with MMEJ-related ID6 indels (B). In contrast, there are several NB tumors with ID6 indels (C). (D-F) Contribution of COSMIC SNV-signature 3 (D) and stacked bar plots of the number of SNVs attributed to COSMIC SNV signatures for Ewing sarcoma tumors (ES) (in E), and neuroblastoma tumors (NB) (in F). Short somatic mutations were determined from whole genome sequencing data of tumors and matched normals using GATK4, and SigProfilerAssignment was used to estimate the COSMIC SNV signatures from the mutations. Each bar represents estimated numbers of SNVs attributed to specific SNV signatures, which are shown in different colors. There are no ES tumors with MMEJ-related Signature 3 SNVs (E). In contrast, there are several NB tumors with Signature 3 SNVs (F). (G-H) Distribution of insertions and deletions by size in Ewing sarcoma tumors (ES; n = 23), and neuroblastoma tumors (NB; n = 47). Short somatic mutations were determined from the whole genome sequencing data of tumors and matched normals. Each bar represents the proportion of deletions (G) or insertions (H) of a particular size among all deletions or all insertions, respectively, of each cohort. (I) Low templated insertions in Ewing sarcoma tumors (ES; n = 23) as compared to neuroblastoma tumors (NB; n = 47). (J, K) The bar graphs show the length of microhomology at canonical translocation breakpoints in Ewing sarcoma samples from GMKF (J, n=46) and St. Jude (K, N=43) cohorts. All translocations were called using the Manta structural variant caller. Canonical translocations are defined as those impacting one of the following genes: EWSR1, FLI1, ETV1, FEV, FUS, ERG.

**Supplementary Figure 6.**
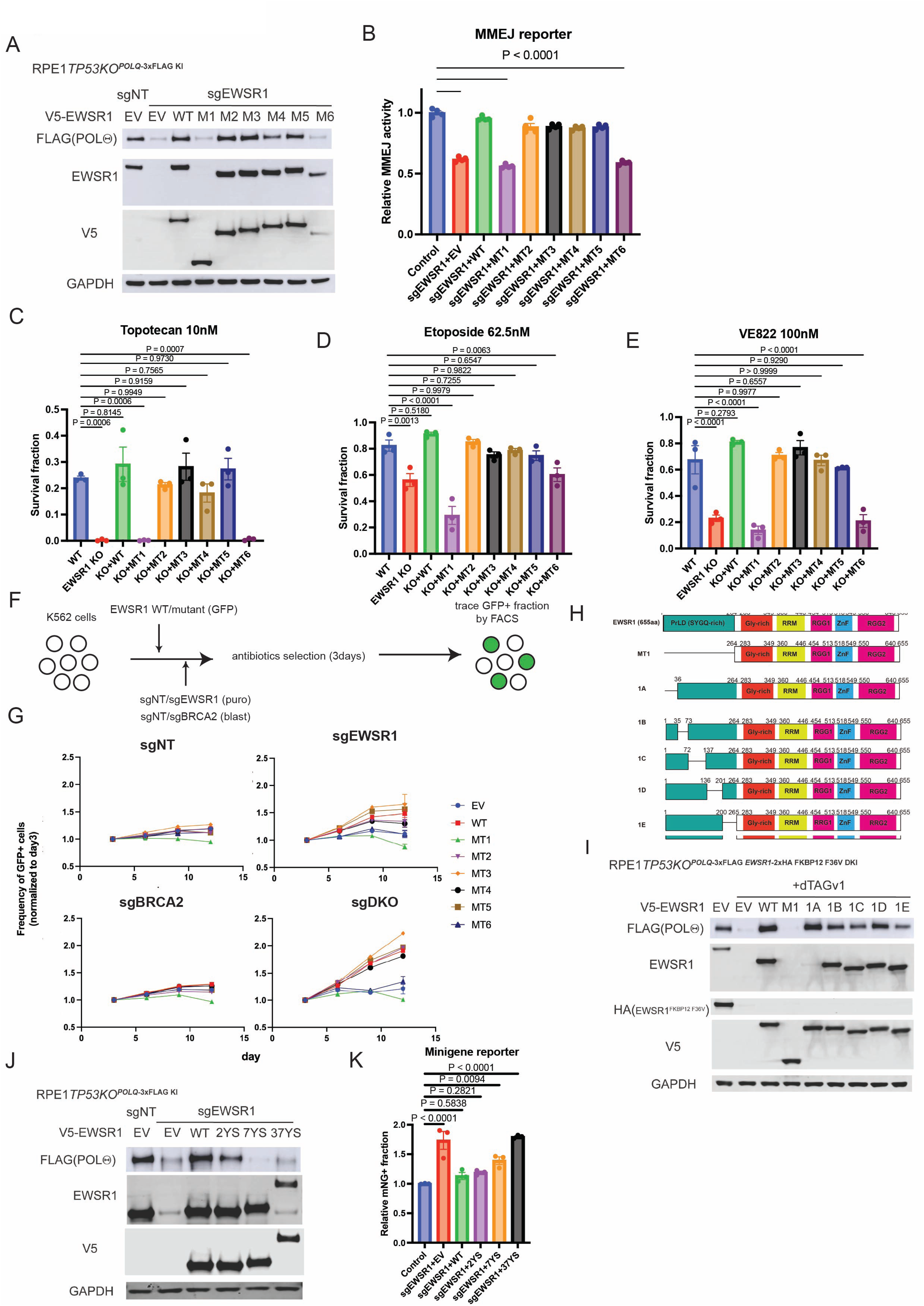
The prion-like domain of EWSR1 is critical for the faithful splicing of *POLQ* transcript. (A) RPE *p53*-/-3xFLAG-*POLQ* knock-in cells were retrovirally transduced with empty vector, wildtype *EWSR1*, or indicated *EWSR1* mutants (co-expressing blasticidin resistance gene) together with lentivirally Cas9 and a control sgRNA or sgRNA targeting *EWSR1* (co-expressing puromycin resistance gene). Cells were treated with puromycin and blasticidin for 4days, and then cell lysates were extracted and then analyzed by immunoblotting. (B) U2OS MMEJ ver.2 (Cas9-based) cells were lentivirally transduced with control sgRNA or sgRNA targeting *EWSR1* together with empty vector wild-type EWSR1 or the indicated EWSR1 mutants by retroviral transduction. Cells were treated with puromycin and blasticidin for 4days and then subjected to MMEJ reporter analysis (n=4). (C-E) Colony formation assay data of RPE *p53*-/-cells and RPE *p53*-/-*EWSR1* knock-out clones transduced with an empty vector, wild-type *EWSR1* or the indicated *EWSR1* mutants by retroviral transduction in the presence of 10nM Topotecan (C), 62.5nM Etoposide (D), or 100nM VE822 (E), respectively. The data is normalized to the DMSO control (n=3). (F, G) K562 cells were lentivirally transduced with sgRNAs targeting control or *EWSR1* (co-expressing puromycin resistance gene) and control or *BRCA2* (co-expressing blasticidin resistance gene). Cells were treated with puromycin and blasticidin for 4days, and then retrovirally transduced with indicated constructs (co-expressing GFP). 72hr after retroviral transduction, the change of frequency of GFP positive fraction was traced by FACS (n=3). (H) Schematic representation of the deletion mutants within the prion-like domain used in Fig. S6I. (I) RPE *p53*-/-3xFLAG-*POLQ*/2xHA-FKBP12 V36F *EWSR1* double knock-in cells were retrovirally transduced with empty vector, wildtype EWSR1, or indicated EWSR1 mutants. After transduction, cells were treated with blasticidin for four days. Then, cells were treated with DMSO or 1μM dTAG^v1^ for 24hr. After dTAG^v1^ treatment, cell lysates were subjected to immunoblot analysis. (J) Wildtype EWSR1 but not the EWSR1 7YS or 37YS mutant rescued POL0 protein expression in RPE *p53*-/-3xFLAG-*POLQ* knock-in cells. (K) Wildtype EWSR1 but not the EWSR1 7YS or 37YS mutant prevented *POLQ* exon 25 skipping measured by K562 minigene reporter cells (n=3).

**Supplementary Figure 7:**
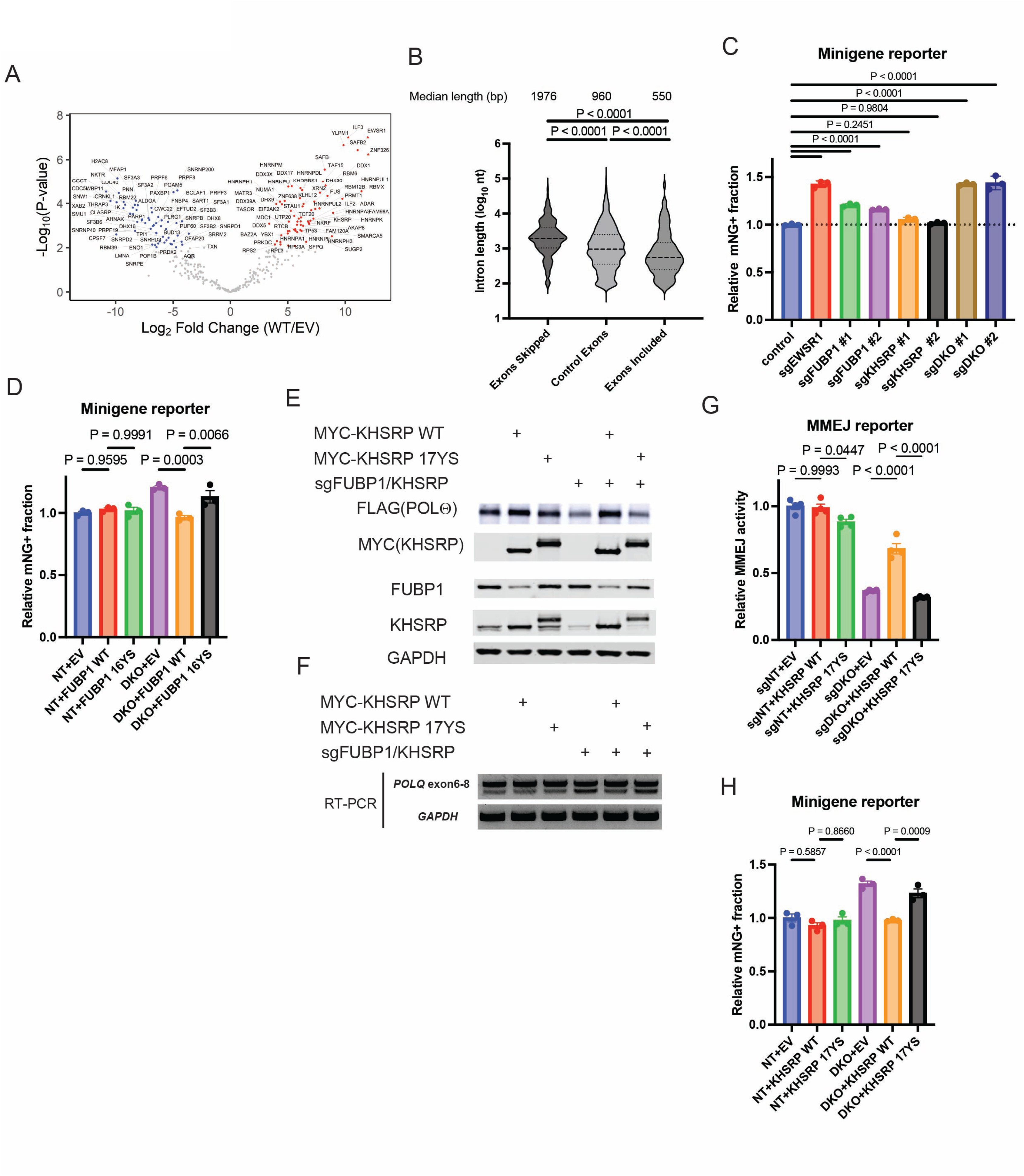
Exon skipping in *EWSR1* KO cells was typically observed for exons that are flanked by longer introns. (A) Volcano plots comparing detected protein levels by IP-MS between EWSR1 wildtype and control. (B) The length of introns surrounding included, excluded or background exons in *EWSR1* knockout cells identified by RNA-seq analysis. (C) Dual depletion of FUBP1 and KHSRP increased *POLQ* exon 25 skipping measured by K562 minigene reporter cells (n=3). (D) The FUBP1 16YS mutant failed to rescue *POLQ* exon 25 skipping induced by FUBP1/KHSRP dual depletion measured by K562 minigene reporter cells (n=3). (E) RPE *p53*-/-3xFLAG-*POLQ*/2xHA-FKBP12 V36F *EWSR1* double knock-in cells were lentivirally transduced with Cas9 and a sgRNA targeting control (NT) or sgRNAs targeting FUBP1 and KHSRP together with retrovirally transduced with a control vector, wildtype KHSRP or the KHSRP 16YS mutant. After puromycin and blasticidin selection, cells were subjected to immunoblotting analysis. (F) RT-PCR showing the *POLQ* exon25 skipping in the indicated cells. (G) U2OS MMEJ ver.2 (Cas9-based) cells were lentivirally transduced with indicated sgRNA(s) (co-expressing puromycin resistance gene). Cells were treated with puromycin for 4days, and then subjected to MMEJ reporter analysis (n=4). (H) The KHSRP 17YS mutant failed to rescue *POLQ* exon 25 skipping induced by FUBP1/KHSRP dual depletion measured by K562 minigene reporter cells (n=3).

**Supplementary Figure 8.**
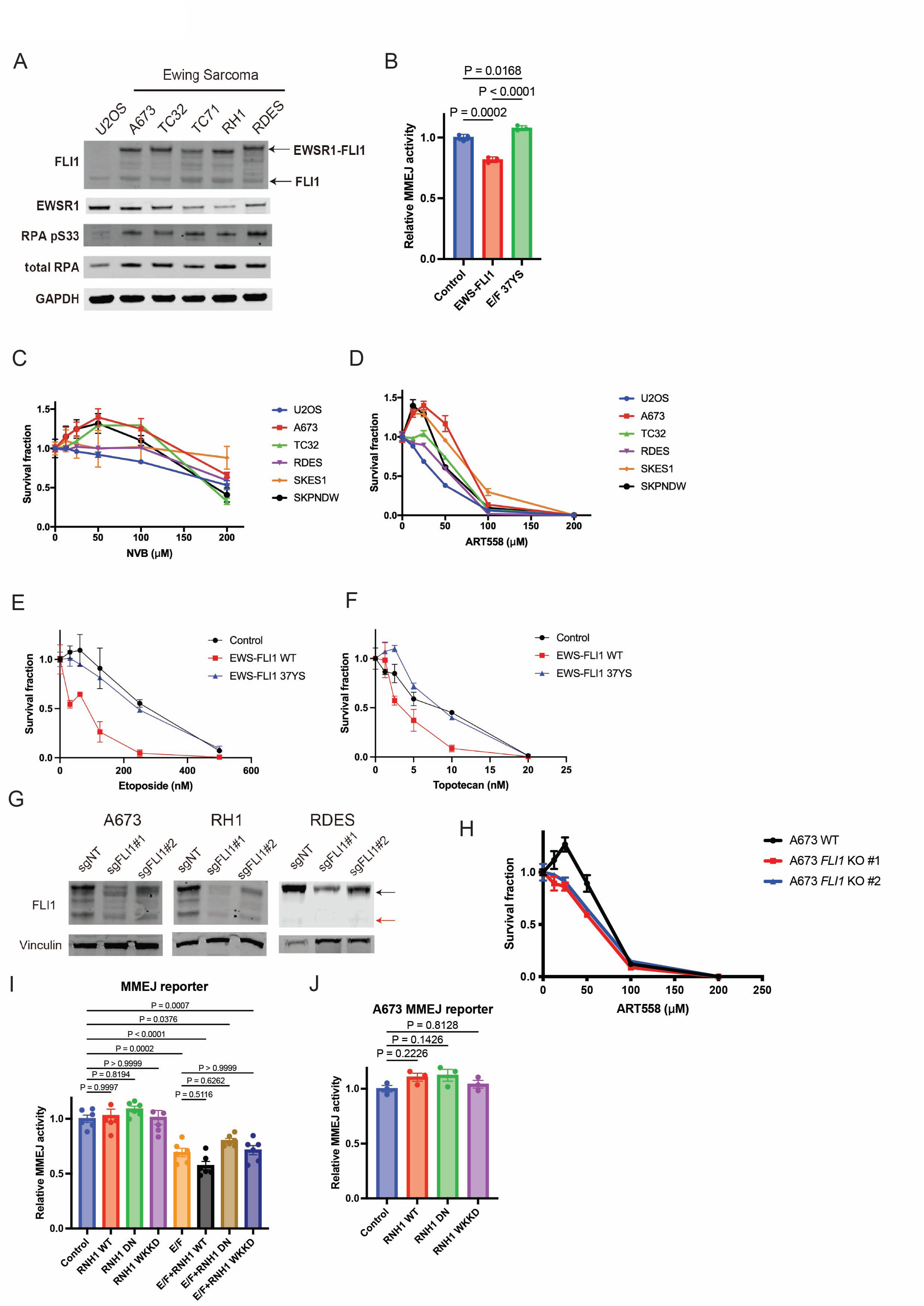
Depletion of EWS-FLI1 ameliorates MMEJ activity in Ewing sarcoma cells. (A) The cell lysates were extracted from an Osteosarcoma cell line U2OS and Ewing Sarcoma cell lines (A673, TC32, TC71, RH1, and RD-ES, and then analyzed by immunoblotting. (B) Cas9-based MMEJ reporter cells were lentivirally transduced with an empty vector, wild-type *EWS-FLI1* or indicated mutant forms of *EWS-FLI1* (co-expressing puromycin resistance gene). After puromycin selection, cells were subjected to MMEJ reporter assay (n=3). (C, D) the CellTiter-Glo® (CTG) assay showing Ewing sarcoma cells showed resistance to POL0 inhibitors, Novobiocin (C) and ART558 (D). (E, F) Colony formation assay data of RPE *p53*-/-cells lentivirally transduced with control vector, EWS-FLI WT or the EWS-FLI1 37YS mutant treated with Etoposide (E) and Topotecan (F) (n=2). (G) The cell lysates used in Fig.4C were extracted and then analyzed by immunoblotting. The black and red arrows showing EWS-FLI1 and FLI1, respectively. (H) The CTG assay shows that depletion of EWS-FLI1 in A673 cells sensitized to ART558. (I) U2OS I-SceI-based MMEJ reporter cells were transduced with control or EWS-FLI1 together with an empty vector, RNaseH1 WT, the RNaseH1 DN, or the RNaseH1 WKKD mutant. Cells were treated with puromycin and blasticidin for 4days and then subjected to MMEJ reporter analysis (n=6). (J) A673 carrying the Isce-I-based MMEJ reporter were transduced with an empty vector, RNaseH1 WT, the RNaseH1 DN, or the RNaseH1 WKKD mutant. Cells were treated with blasticidin for 4days and then subjected to MMEJ reporter analysis (n=3).

**Supplementary Figure 9.**
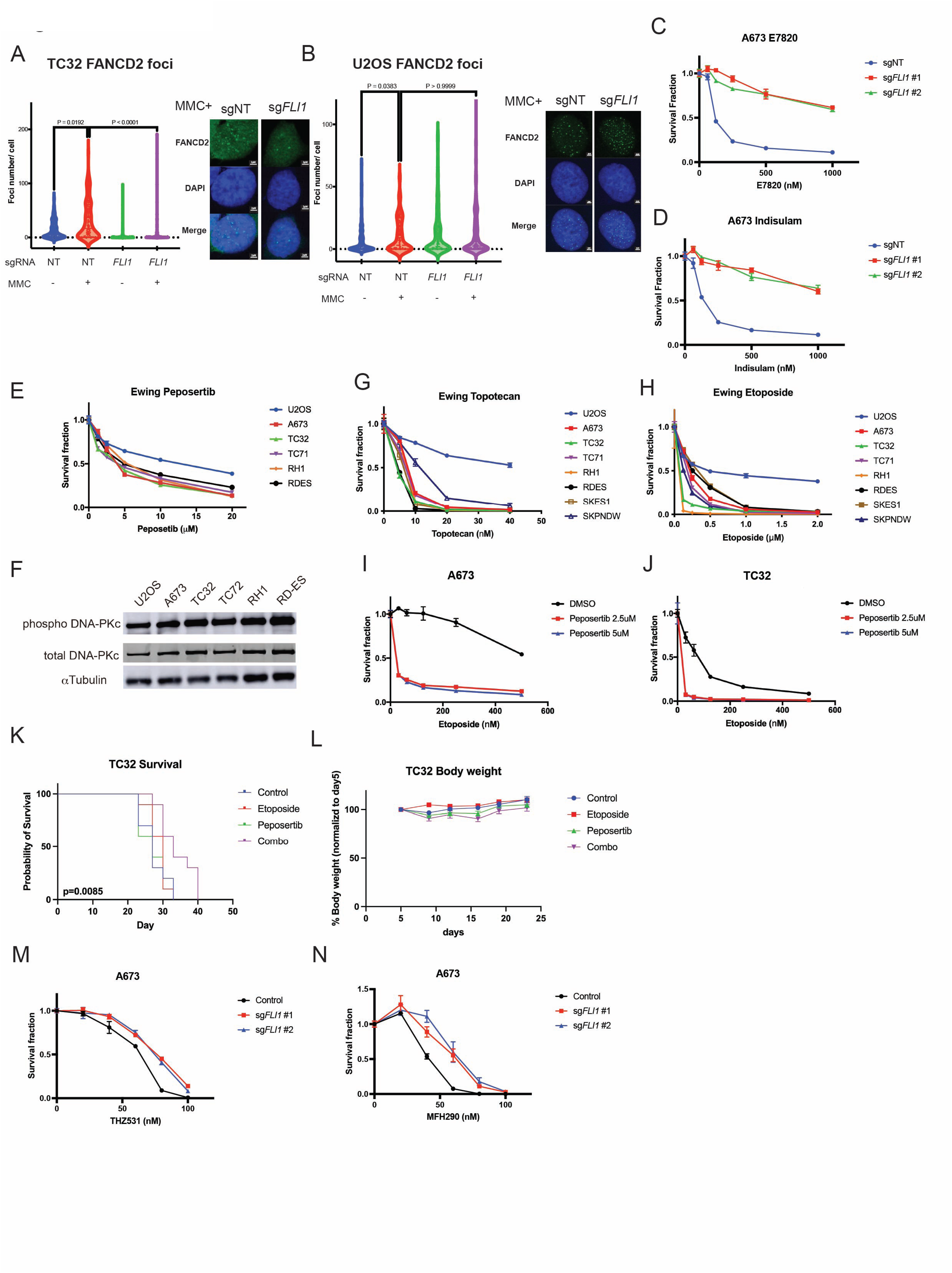
Combination of Peposertib and Etoposide efficiently killed Ewing sarcoma cells in vivo. (A, B) Immunofluorescence analysie showing FANCD2 foci formation was increased by EWS-FLI1 expression in TC32 cells (A) but not by sgFLI1 in U2OS cells (B). Cells were treated with 10ng/ml MMC for 24hr. Shown are the number of FANCD2 foci (left) and the representative images of the cells treated with MMC (right, scale bars = 2um). (C, D) The CTG assay showing that depletion of EWS-FLI1 by sg*FLI1* in A673 cells led to acquisition of resistance to RBM39 degraders, E7820 (C) and Indisulam (D). (E) The CTG assay showing that compared with U2OS, Ewing sarcoma cells showed hypersensitivity to Peposertib. (F) Immunoblot analysis showing phospho-DNAPK, DNAPK and GAPDH protein level in U2OS and Ewing sarcoma cell lines. (G, H) The CTG assay showing that compared with U2OS, Ewing sarcoma cells showed hypersensitivity to Topotecan (G) and Etoposide (H). (I, J) The CTG assay showing that a combination of Peposertib and Etoposide showed a strong synergistic effect on Ewing sarcoma cell lines A673 (I) and TC32 (J). (K) Overall survivals showing a therapeutic efficacy of a combination of Etoposide and Peposertib in a TC32 xenograft model (n=10, each group). (L) The changes of the body weight of the treated mice. (M, N) The CTG assay showing that depletion of EWS-FLI1 in A673 cells led to acquisition of resistance to CDK12 inhibitors, THZ531 (M) and MFH290 (N).

**Supplementary Figure 10.**
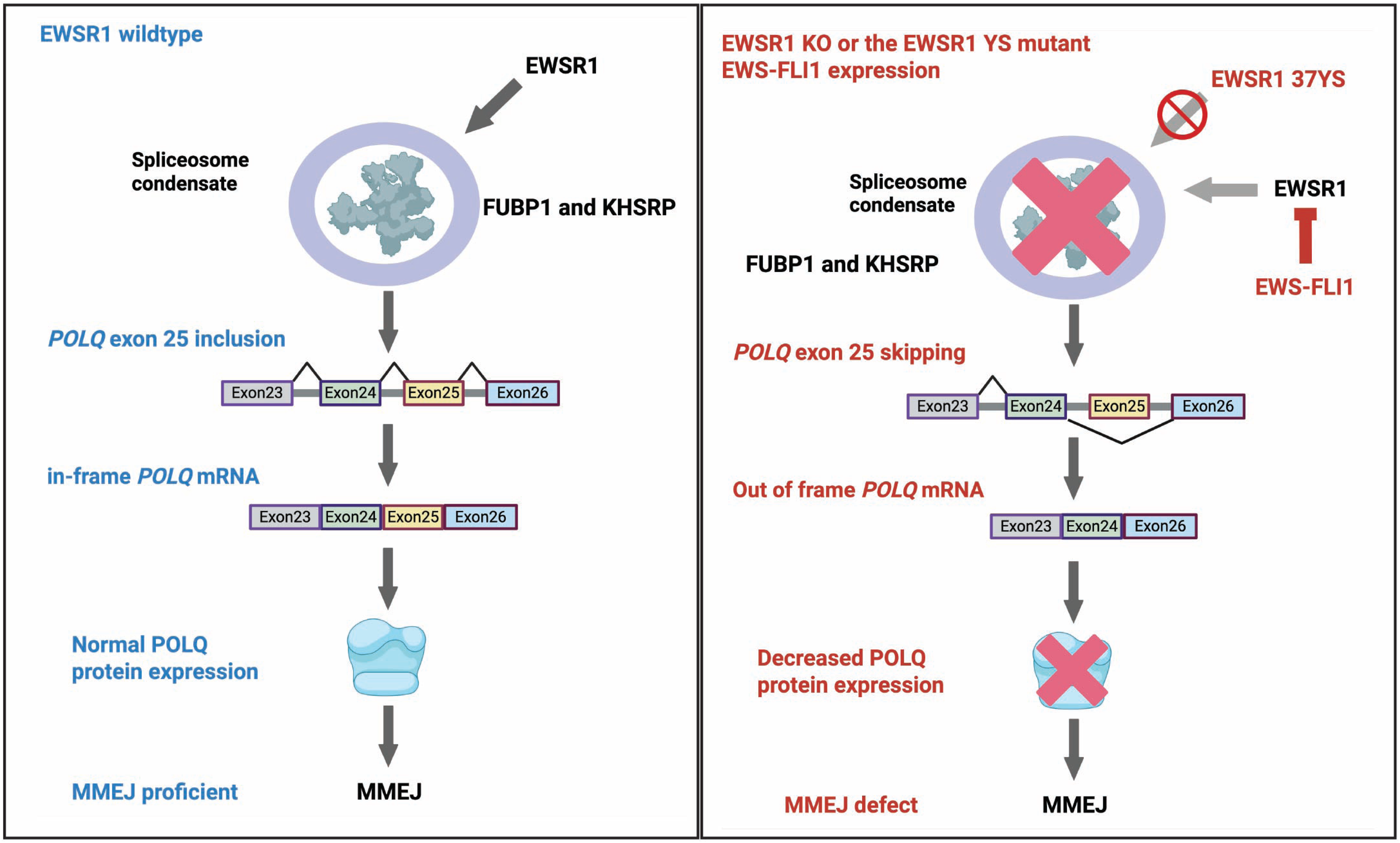
The schematic representation of the findings of this study.

